# Unstructured mRNAs form multivalent RNA-RNA interactions to generate TIS granule networks

**DOI:** 10.1101/2020.02.14.949503

**Authors:** Weirui Ma, Gang Zhen, Wei Xie, Christine Mayr

## Abstract

The TIS granule network is a constitutively expressed membraneless organelle that concentrates mRNAs with AU-rich elements and interacts with the major site of protein synthesis, the rough endoplasmic reticulum. Most known biomolecular condensates are sphere-like, but TIS granules have a mesh-like morphology. Through *in vivo* and *in vitro* reconstitution experiments we discovered that this shape is generated by extensive intermolecular RNA-RNA interactions. They are mostly accomplished by mRNAs with large unstructured regions in their 3′UTRs that we call intrinsically disordered regions (IDRs). As AU-rich RNA is a potent chaperone that suppresses protein aggregation and is overrepresented in mRNAs with IDRs, our data suggests that TIS granules concentrate mRNAs that assist protein folding. In addition, the proximity of translating mRNAs in TIS granule networks may enable co-translational protein complex formation.

---

Biomolecular condensates form through weak interactions of multivalent molecules. Protein-protein interactions occur between repeated modular domains or between intrinsically disordered regions (IDRs). Protein IDRs lack a defined three-dimensional structure but often contain low-complexity sequence elements that provide the basis for multivalent intermolecular interactions (*1*). In addition, protein-RNA and RNA-RNA interactions contribute to the multivalency of phase separation systems (*2*-*7*). For example, it has been demonstrated that RNA can phase separate without protein and that RNA can promote or inhibit phase separation (*3, 8*). It has also been shown that protein-RNA interactions can influence the identity and material properties of condensates *in vitro* and *in vivo* (*2, 4, 6*). However, the contribution of specific mRNAs to phase separation is largely unknown, as it has only been studied for few mRNAs (*2, 4*).

The TIS granule network is a membraneless organelle that forms a mesh-like compartment that is intertwined with the endoplasmic reticulum (ER). It is present in all cell types studied under physiological conditions. For mRNA to localize to TIS granules, they require several AU-rich elements in their 3′ untranslated regions (3′UTRs) (*9*). Whereas most known biomolecular condensates are sphere-like (*1*), TIS granules form a tubule-like meshwork. Here, we set out to investigate how the characteristic three-dimensional organization of TIS granules is determined.

During these studies, we examined the influence of 47 human *in vitro* transcribed 3′UTRs, with sizes ranging from 500-5000 nucleotides, on phase separation of an RNA-binding protein. We observed that mRNAs that form strong local intramolecular interactions generate sphere-like condensates. In contrast, extensive intermolecular RNA-RNA interactions drive the generation of mesh-like condensates and are accomplished by mRNAs with large unstructured regions that are often single-stranded and AU-rich. Taken together, the TIS granule network represents a cytoplasmic compartment that concentrates unstructured, AU-rich mRNAs with a high propensity for intermolecular interactions. Intermolecular mRNA-mRNA interactions during translation may provide proximity of nascent chains to allow co-translational assembly of protein complexes. Intermolecular mRNA-protein interactions in TIS granules may enable chaperone function of mRNA to assist in protein folding.

TIS granules form through assembly of the RNA-binding protein TIS11B. Expression of GFP-tagged TIS11B in HeLa cells recapitulates the mesh-like three-dimensional structure of endogenous TIS granules (Fig. 1A) (*9*). TIS11B binds to AU-rich elements through a double zinc finger RNA-binding domain. Introduction of different point mutations to disrupt RNA binding (fig. S1, A and B) (*10*) turned the mesh-like assemblies into sphere-like condensates that are no longer intertwined with the ER (Fig. 1A, and fig. S1, C and D). This suggested that the mesh-like organization of TIS granules requires either an intact zinc finger domain or is caused by the bound mRNAs. To distinguish between the two possibilities, we generated a chimeric protein where we replaced the RNA-binding domain of TIS11B with RRM1/2 from HuR, as both RNA-binding domains bind to AU-rich elements in 3′UTRs (fig. S1B) (*11*). As this chimeric protein retained the ability to form a granule network (Fig. 1B), the data suggests that mRNAs bound by either HuR or TIS11B are responsible for the mesh-like organization of TIS granules.

**Figure 1.**
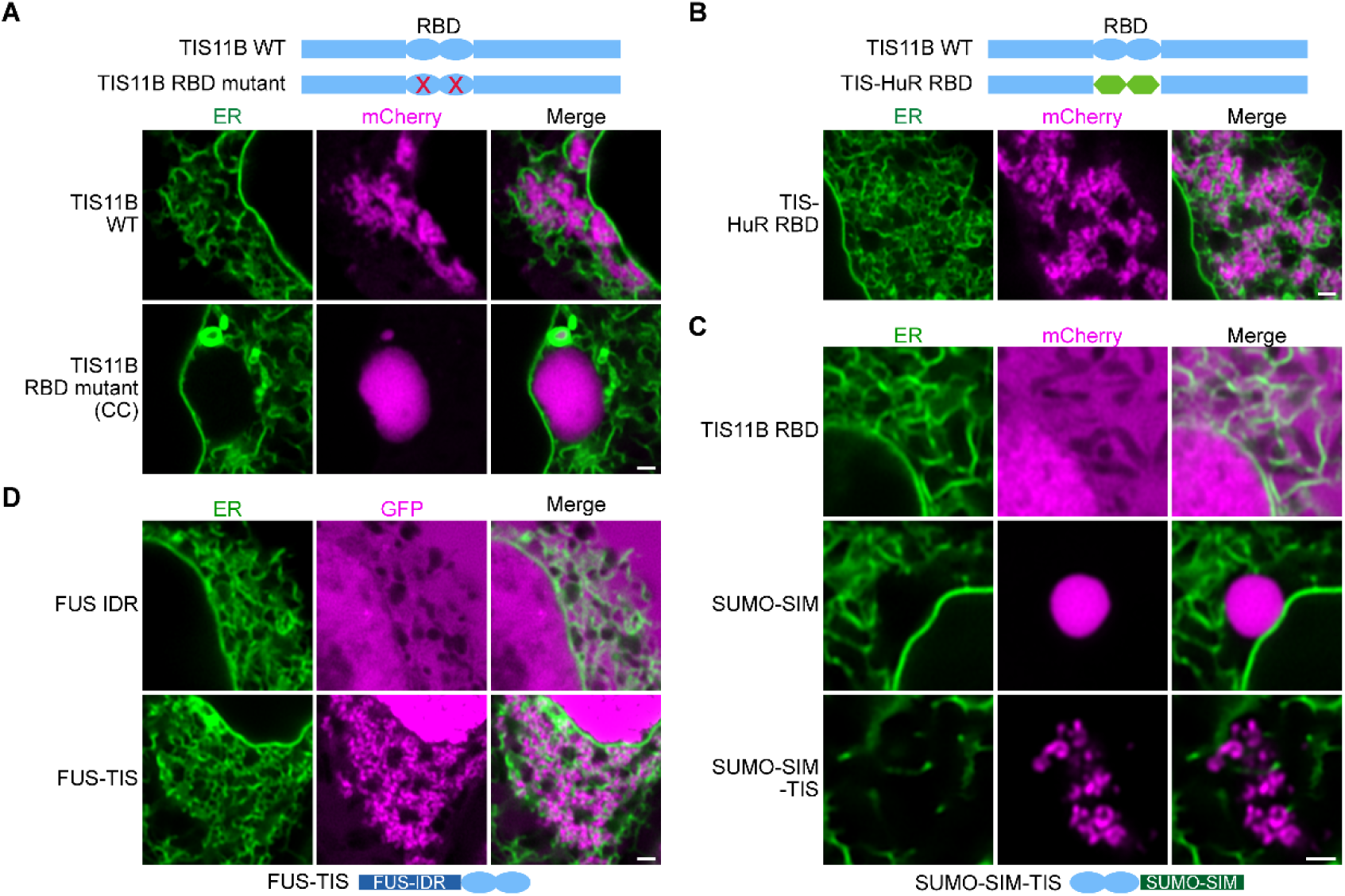
RNA-binding determines the mesh-like three-dimensional organization of TIS granules in cells. **(A)** Confocal live cell imaging of HeLa cells after the transfection of mCherry-tagged TIS11B orTIS11B with mutated RNA-binding domain (RBD). GFP-SEC61B was co-transfected to visualize the ER. Scale bar, 1 μm; CC, C135H/C173H. See fig. S1 for more mutants. **(B)** Same as (A), but after transfection of TIS-HuR RBD chimera, All granules are mesh-like. **(C)** Same as (A), but after transfection of TIS11B RBD, SUMO10-SIM5 or SUMO-SIM-TIS chimera. 73% (*N* = 52) of SUMO-SIM-TIS granules are mesh-like. **(D)** Same as (A), but after transfection of mGFP-tagged FUS IDR (amino acids 1-214) or FUS-TIS chimera. All granules are mesh-like. mCherry-SEC61B was co-transfected to visualize the ER.

Expression of the TIS11B RNA-binding domain alone does not result in condensate formation (Fig. 1C). We investigated if the TIS11B RNA-binding domain is sufficient for network formation in the context of different multivalent domains and fused it to SUMO-SIM (SUMO-SIM-TIS) or to the IDR of FUS (FUS-TIS) (*12, 13*). Both chimeric proteins form granule networks *in vivo* (Fig. 1, C and D, fig. S1E). As SUMO-SIM-TIS creates a tubule-like condensate that is not intertwined with the ER, this experiment revealed that intertwinement with the ER is not necessary for mesh-like network formation of TIS chimeras (Fig. 1C, fig. S1E). These data indicate that the TIS11B RNA-binding domain drives granule network formation in the context of diverse multivalent domains.

To learn the determinants of mRNAs that enable network formation, we turned to an *in vitro* approach. Due to aggregation, we were not able to purify monomeric TIS11B. Instead, we recombinantly expressed and purified FUS-TIS (fig. S2, A and B), as it forms granule networks in cells. FUS-TIS phase separates into liquid-like and sphere-like condensates (fig. S2, C and D). We then added to FUS-TIS *in vitro* transcribed 3′UTRs of mRNAs that were previously shown to localize to TIS granules (*9*). Whereas the addition of the *FUS* 3′UTR did not change the morphology of the sphere-like condensates formed by FUS-TIS, the addition of different 3′UTRs obtained from *CD47, CD274* (PD-L1) or *ELAVL1* (HuR) resulted in the formation of mesh-like FUS-TIS condensates (Fig. 2A, fig. S3). The mesh-like condensates do not represent aggregates as FUS-TIS protein showed fast fluorescence recovery after photobleaching (Fig. 2B).

**Figure 2.**
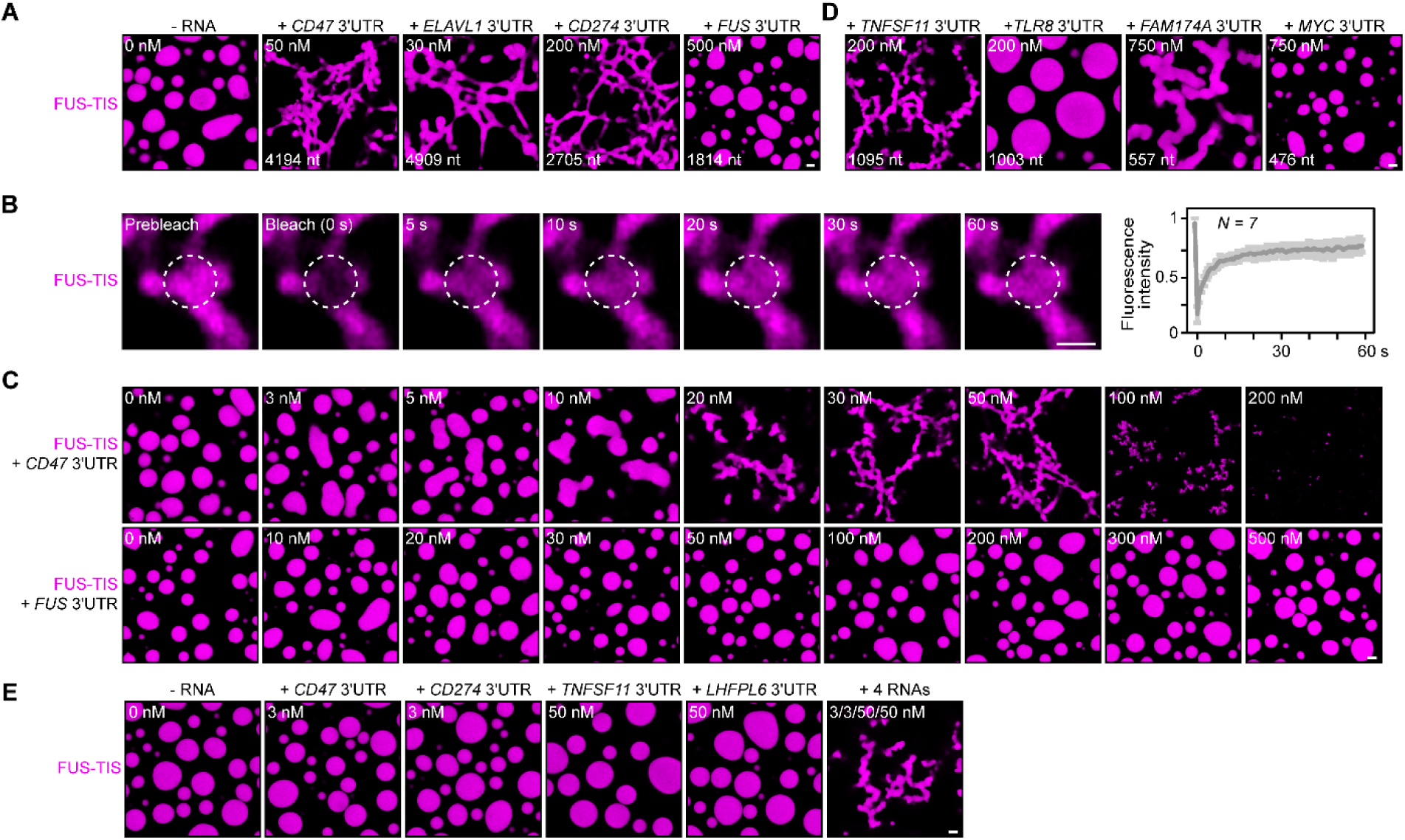
Specific RNAs induce granule network formation *in vitro*. **(A)** Representative confocal images of phase separation experiments using purified mGFP-FUS-TIS (10 μM) in the absence or presence of the indicated in vitro transcribed RNAs after 16 hours of incubation. Scale bar, 2 μm. **(B)** Fluorescence recovery after photobleaching of mGFP-FUS-TIS (10 μM) mixed with *CD47* 3′UTR (50 nM) after two hours of incubation. Scale bar, 1 μm. **(C)** Same as (A), but in the presence of different RNA concentrations. **(D)** Same as (A), but additional RNAs are shown. **(E)** Representative confocal images of phase separation experiments using purified mGFP-FUS-TIS (10 μM) in the presence of a single network-forming RNA at sub-optimal concentration, or in the presence of four network-forming RNAs, each at sub-optimal concentration. Images were taken after 16 hours of incubation.

All phase separation experiments were performed at two timepoints (after two and 16 hours of incubation) and at RNA concentrations spanning three orders of magnitude. Network formation was already observed at the early time point, but longer incubation led to formation of a more extensive network (Fig. 2A, fig. S3, S4). Although the minimum RNA concentration required to induce network formation varied, these experiments revealed that the capacity for network formation is an intrinsic property of the RNA, as sphere-forming RNAs did not form networks even at high concentration. Instead, at high RNA concentrations, we often observed inhibition of phase separation, as was observed previously (Fig. 2C, fig. S3) (*8*).

The three network-forming RNAs are longer than the sphere-forming RNA (Fig. 2A). To examine if network formation is only accomplished by long RNAs, we tested 19 additional RNAs with a length spanning 500 – 3000 nt. All longer RNAs formed networks, but we observed both network and sphere formation for RNAs shorter than 2000 nt, indicating that network formation is not only determined by the length of the RNA (fig. S5 and S6).

The minimum RNA concentration for network formation was 20 nM. It was observed for the *CD47* and *ELAVL1* 3′UTRs and corresponds to 27 and 32 ng/μl, respectively (Fig. 2C and fig. S3). This is higher than the mRNA concentration in the cytoplasm of mammalian cells which was estimated to be 8 pM to 8 nM (9.5 pg/μl to 9.5 ng/μl; see methods) (*8, 14, 15*). As TIS granules contain many mRNAs (*9*), we hypothesized that multiple RNAs may contribute to network formation. Therefore, we tested whether two RNAs co-localize in the network. Labeling of several RNAs with two different fluorescent dyes showed that they co-localize (fig. S7). Furthermore, the mixing of sub-optimal amounts of several network-forming RNAs together with FUS-TIS resulted in network formation, indicating that the different RNAs have an additive effect (Fig. 2E). As thousands of mRNAs contain AU-rich elements in their 3′UTRs (table S2), our data suggests that many different mRNAs may contribute to the formation of TIS granule networks in cells.

As network formation is an intrinsic feature of certain RNAs, we set out to identify the responsible determinants. We used 18 different 3′UTRs that were shorter than 2000 nt and correlated the ability to form networks with several parameters. Within this size-restricted cohort, the number of AU-rich elements or the GC-content of the RNA had no influence on network formation (fig. S8, A and B). We then used RNAfold to predict the secondary structure of the RNAs and found that RNAs with large regions of strong local structure, indicated by a high base-pairing probability (red color code), were unable to form networks (Fig. 3A, fig. S8, C and D, and table S1) (*16*). In contrast, RNAs that contained large unstructured regions, indicated by the green and blue colors, had a high propensity for network formation (Fig. 3B, fig. S8, E and F, and table S1). We call the unstructured regions ‘intrinsically disordered regions’ (IDRs) of mRNAs.

**Figure 3.**
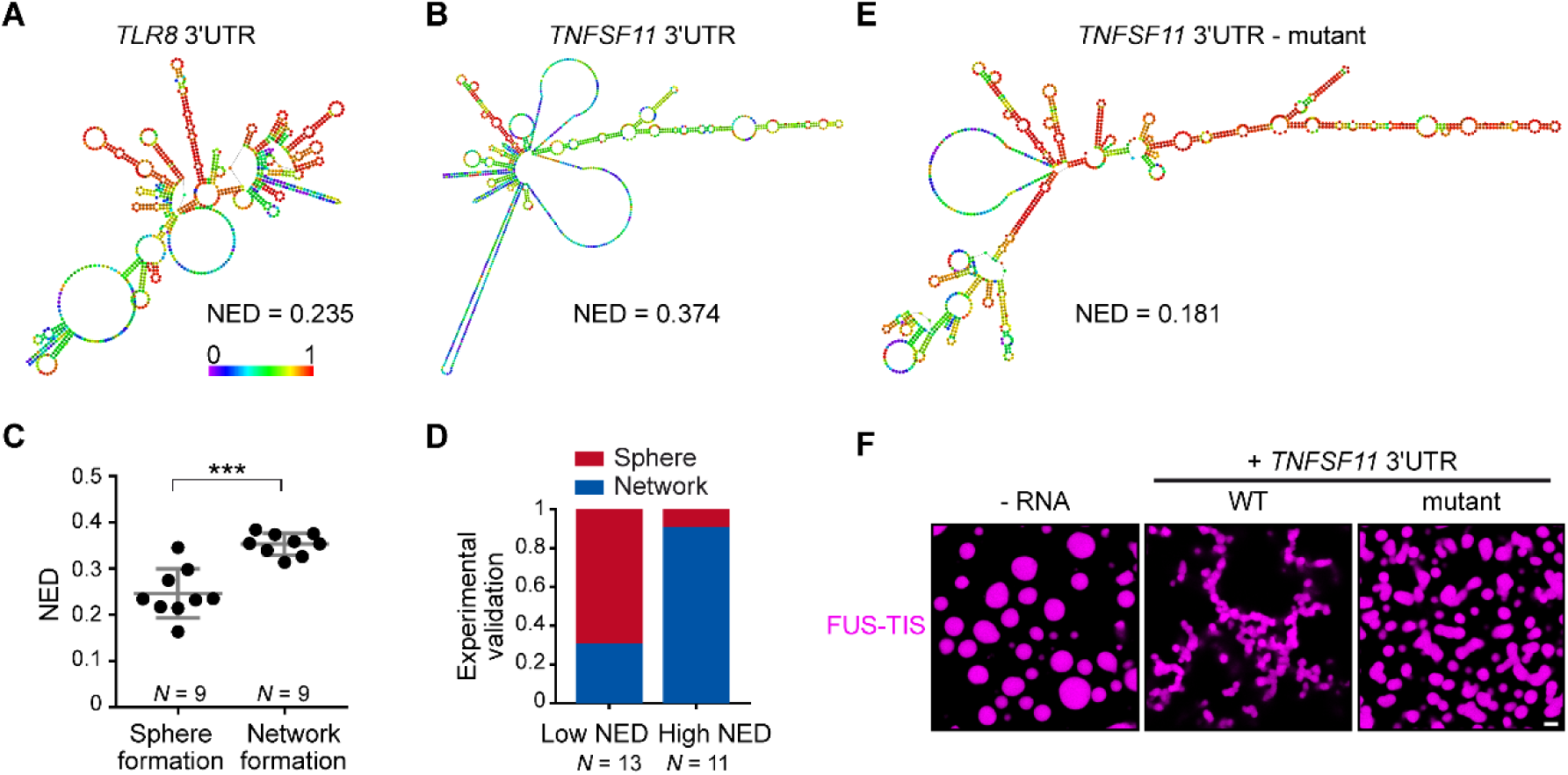
RNAs with large disordered regions have a high propensity for network formation. **(A)** Centroid RNA secondary structure of *TLR8* 3′UTR predicted by RNAfold. The color code represents base-pairing probability. **(B)** Same as (A), but the *TNFSF11* 3′UTR is shown. **(C)** Normalized ensemble diversity (NED) values of sphere- and network-forming RNAs. See table S1. Mann-Whitney test, Z=-3.3, ***, *P* < 0.0003. **(D)** Experimental validation of *N* = 24 in vitro transcribed RNAs whose ability for network formation was predicted by NED. Sphere formation is indicated in red, whereas network formation is indicated in blue. See table S1. Mann-Whitney test was performed on the experimental validation, Z=-2.8, ***, *P* = 0.004. **(E)** Same as (B), but the mutant *TNFSF11* 3′UTR is shown. **(F)** Representative confocal images of phase separation experiments using purified mGFP-FUS-TIS (10 μM) in the presence of 150 nM of the indicated in vitro transcribed RNAs after 16 hours of incubation. Scale bar, 2 μm.

While trying to assign numeric values to the base-pairing probability color code, we noticed that the parameter of ‘ensemble diversity’ correlates strongly with the ability for network formation. Ensemble diversity is the number of potential RNA structures that are predicted for a given RNA (*17*). As this value increases with RNA length (*18*), we are using a length-normalized value (NED). We observed that RNAs that form strong local structures have low NED values, whereas RNAs with IDRs have high values (Fig. 3C). Intriguingly, the NED values separated the two groups of RNAs with respect to network formation.

To test the predictive value of NED, we chose a new set of 24 3′UTRs purely based on their NED values and tested their network-forming abilities. We found that 19/24 (79%) of the tested RNAs were predicted correctly with respect to their sphere- or network-forming ability (Fig. 3D, fig. S9 and S10, and table S1). The high success rate strongly suggests that IDRs of 3′UTRs determine network formation. To test this prediction experimentally, we performed a loss-of-function experiment. We used the *TNFSF11* 3′UTR that contains large IDRs (Fig. 3B) and introduced strong local base-pairing by the addition of two oligonucleotides that were perfectly complementary to upstream regions (fig. S11). The *TNFSF11* 3′UTR mutant has stronger local structures indicated by increased base-pairing and lower NED values, and it has largely lost the ability for network formation (Fig. 3, E and F). Taken together, these results support a model wherein a high diversity of predicted structural conformations correlates with the extent of IDRs in RNAs and is associated with formation of mesh-like condensates.

Next, we set out to address how RNAs with IDRs form networks. We had observed that RNAs that are unable to form networks are predicted to form strong local structures, meaning that they have a high propensity for intramolecular interactions (Fig. 4A). This led us to hypothesize that network formation is caused by intermolecular RNA-RNA interactions mediated by the IDRs of mRNAs (Fig. 4B). To test this, we performed native gel electrophoresis with sphere-forming and network-forming RNAs. We observed the appearance of diverse RNA species with high molecular weight only with the network-forming RNAs (Fig. 4C). This suggests that mRNAs with IDRs form higher-order RNA interactions *in vitro*.

**Figure 4.**
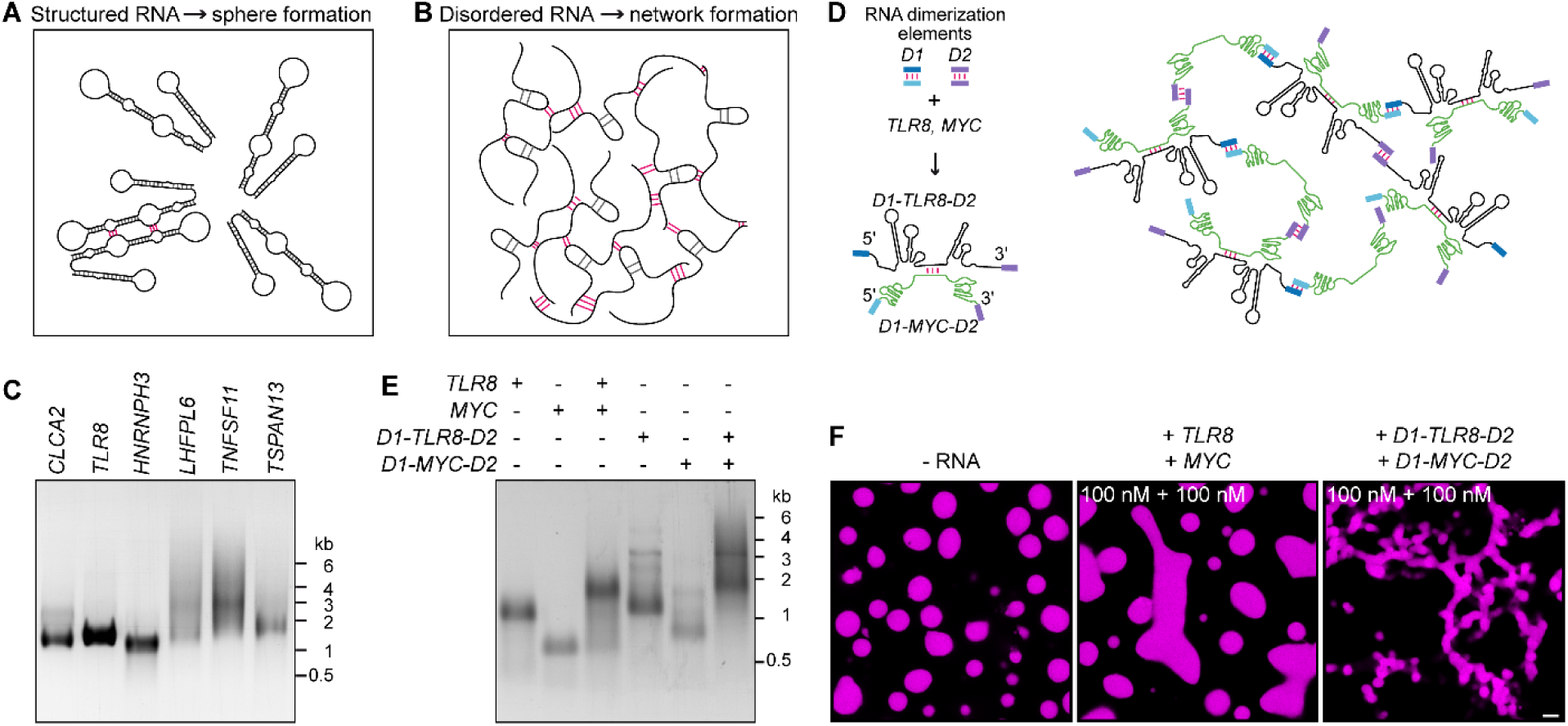
Extensive intermolecular RNA-RNA interactions are responsible for granule network formation. **(A)** Schematic for RNAs with strong local secondary structures. **(B)** Schematic for RNAs with large IDRs that form extensive intermolecular RNA-RNA interactions. **(C)** Native agarose gel electrophoresis of sphere-forming (lanes 1-3) and network-forming (lanes 4-6) RNAs (5 μM, each). **(D)** Schematic of complex RNA network characterized by extensive intermolecular RNA-RNA interactions mediated by different dimerization elements (*D1* and *D2*) that were added to structured RNAs. **(E)** Native agarose gel electrophoresis of the indicated RNAs (1 μM, each). **(F)** Representative images of phase separation experiments using purified FUS-TIS (10 μM) in the presence of the indicated *in vitro* transcribed RNAs after 16 hours of incubation. Scale bar, 2 μm.

To investigate if granule networks are indeed caused by intermolecular RNA-RNA interactions, we set out to reconstitute a mesh-like condensate using RNAs that were designed to form multivalent RNA-RNA interactions (Fig. 4D). We selected two 3′UTRs that are unable to induce network formation (Fig. 2D), but that are able to dimerize, thus, providing one degree of multivalency (Fig. 4, D and E, fig. S12). We added two different RNA dimerization elements to their 5′ and 3′ ends to increase RNA multivalency. This strategy enables intermolecular RNA-RNA interactions and allows the formation of a complex RNA network, demonstrated by native gel electrophoresis (Fig. 4, D and E). The RNA dimerization elements were derived from tracrRNA/crRNA (*D1*) and from HIV (*D2*; fig. S13, A and B) (*19, 20*).

A phase separation experiment with FUS-TIS confirmed that the addition of the two predominantly structured 3′UTRs (*TLR8* and *MYC*) generates sphere-like condensates, whereas the addition of RNAs capable of forming extensive intermolecular RNA-RNA interactions (*D1-TLR8-D2* and *D1-MYC-D2*) induces formation of a mesh-like FUS-TIS condensate (Fig. 4F and fig. S13C). Taken together, this *in vitro* reconstitution experiment demonstrated that mesh-like condensates are caused by extensive intermolecular RNA-RNA interactions and high RNA multivalency.

All mRNAs have unstructured and structured regions (*21*). Here, we found that 3′UTRs with large regions of strong base-pairing induce formation of sphere-like condensates, whereas 3′UTRs with large single-stranded, unstructured regions result in network-like condensates that are accomplished by multivalent RNA-RNA interactions. This raises the question about the purpose of having large unstructured regions in mRNA.

To better understand the functions of mRNAs with large IDRs, we performed transcriptome-wide analyses on the co-occurrence of IDRs with specific mRNA features. In HeLa cells, approximately a third of mRNAs (32.5%) have high NED values, suggesting that they have large IDRs in their 3′UTRs (table S2, fig. S14A). As TIS granules enrich mRNAs with several AU-rich elements (*9*), we examined if the number of AU-rich elements correlates with the ensemble diversity parameter of 3′UTRs and found a very strong correlation (fig. S14B). As both features correlate with 3′UTR length, we tested their association after using length-normalized NED values. We still found a strong positive correlation between AU-rich elements and NED (Fig. S14C). This indicates that mRNAs that are enriched in TIS granules are enriched in IDRs. As the mesh-like three-dimensional organization of TIS granules depends on the presence of specific RNAs *in vivo*, we conclude that the TIS granule network represents a cytoplasmic compartment that concentrates mRNAs with AU-rich elements and IDRs. Our data indicates that one function of unstructured mRNAs is to allow multivalent RNA-RNA interactions which is the basis of mesh-like condensates. Therefore, unstructured mRNAs that are enriched in TIS granules are required for its intertwinement with the ER (*9*).

Another consequence of the pervasive intermolecular RNA-RNA interactions that happen in TIS granules may be to provide proximity of two translating ribosomes on different messages. It was shown in yeast that the majority of protein complexes form co-translationally, however, currently, it is unclear how the protein subunits come into proximity (*22, 23*). Physical interaction of the IDRs in the 3′UTRs would bring their nascent chains close together and may promote co-translational protein complex formation.

We then noticed that mRNAs with IDRs generally have a higher AU-content and a predominant enrichment of uridines (fig. S14D). These features were substantially more enriched when we focused on mRNAs containing both AU-rich elements (>3) and IDRs (fig. S14E). Intriguingly, these mRNAs preferentially encode large proteins as well as proteins with large IDRs (fig. S14, F and G). We previously showed that the TIS granule network localizes to the surface of the rough ER, which is the primary site of protein synthesis in most cells (*9*). As large proteins are more difficult to fold and proteins with large IDRs have a tendency to aggregate (*24, 25*), we propose that translation in TIS granules provides a favorable environment during translation and folding of ‘difficult to express’ proteins under physiological conditions. This is based on the recent finding that RNA has a very potent chaperone function *in vitro*. It was demonstrated that RNA is more effective in the suppression of aggregation than conventional chaperones and that RNA can cooperate with protein chaperones during protein refolding (*26, 27*). This function of RNA is further supported by the observation that a high RNA concentration in the nucleus solubilizes proteins with prion-like domains (*8*). Moreover, we have found previously that protein chaperones, including HSC70 are enriched in the TIS granule network (*9*). Here, we found that TIS granules concentrate mRNAs with single-stranded, unstructured regions with a high AU-content. These features are the characteristics of RNAs with chaperone function, as was shown before (*26, 27*).

We propose that unstructured mRNA does not only have a propensity for mRNA-mRNA interactions, but also for mRNA-protein interactions with a preference for unfolded and disordered proteins. This would explain previous observations that found that protein IDRs account for half of all RNA binding events (*28, 29*). The increased affinity for unfolded proteins would also explain the function of RNA as protein chaperones during translation. In the future, it will be important to learn how RNA assists protein folding and prevents protein aggregation.

## Acknowledgements

We thank all members of the Mayr lab for helpful discussions and critical reading of the manuscript. We thank Juncheng Wang for suggestions for protein purification. We also thank Bede Portz and James Shorter for attempting the purification of TIS11B and for valuable suggestions on phase separation experiments. This work was funded by the NIH Director′s Pioneer Award (DP1-GM123454), the Pershing Square Sohn Cancer Research Alliance, and the NCI Cancer Center Support Grant (P30 CA008748). W.M. performed all experiments and analyses. W.X. provided help with protein purification. G.Z. calculated the NED values, identified disordered proteins, and performed the transcriptome-wide analyses. W.M. and C.M. conceived the project, designed the experiments, and wrote the manuscript.

The authors declare no competing interests.

## Materials and Methods

### Cell lines

The human cervical cancer cell line, HeLa, was a gift from the lab of Jonathan S. Weissman (UCSF), provided by Calvin H. Jan. Cells were maintained at 37°C with 5% CO2 in Dulbecco’s Modified Eagle Medium (DMEM) containing 4,500 mg/L glucose, 10% heat-inactivated fetal bovine serum, 100 U/ml penicillin, and 100 mg/ml streptomycin.

### Constructs

All primers are reported in table S3. All PCR reactions were performed using Q5 High Fidelity DNA polymerase (NEB). The basis for all mammalian expression vectors was pc-DNA-puro described previously (*9*). The following inserts were also described previously mCherry-TIS11B, mCherry-SEC61B, eGFP-SEC61B, eGFP-CD47-3′UTR, eGFP-ELAVL1-3′UTR, eGFP-FUS-3′UTR, and eGFP-CD274-3′UTR (*9*).

Point mutations were generated using QuikChange Lightning Multi Site-Directed Mutagenesis Kit (Agilent Technologies, #210513) if not otherwise stated. pcDNA-puro-mGFP (monomeric GFP, A207K) was generated from pcDNA-puro-eGFP using the primer eGFP A207K. For TIS11B RBD (RNA-binding domain) mutants the following primers were used: TIS11B C135H, TIS11B C173H, TIS11B F137N, TIS11B F175N, TIS11B K116L, TIS11B K154L, TIS11B R116L, and TIS11B K152L.

To generate TIS-HuR RBD (TIS11B contains the HuR RBD instead of TIS11B RBD) chimera, three overlapping fragments were PCR amplified. Fragment 1: the N-terminus of TIS11B (aa 1 to 113, based on uniprot ID Q07352-1), was PCR amplified from the mCherry-TIS11B construct with primers TIS-HuR 1F and TIS-HuR 1R. Fragment 2, the RRM1/2 (aa 19 to 189) of HuR, was PCR amplified from the eGFP-HuR-3′UTR construct with primers TIS-HuR 2F and TIS-HuR 2R. Fragment 3: the C-terminus of TIS11B (aa 182 to 338), was PCR amplified from the mCherry-TIS11B construct with primers TIS-HuR 3F and TIS-HuR 3R. A ligation PCR was performed to ligate Fragment 1 and Fragment 2 to generate Fragment 1-2 with primers TIS-HuR 1F and TIS-HuR 2R-2. Then Fragment 1-2 was digested with HindIII and ApaI; Fragment 3 was digested with ApaI and EcoRI. To generate full length TIS-HuR chimera, Fragment 1-2 and Fragment 3 were cloned into the pcDNA3.1-puro-mCherry vector with HindIII and EcoRI restriction sites.

The pmCherry-SUMO10-SIM5 construct was a gift from the lab of Liam J. Holt (NYU). To generate the SUMO-SIM-TIS (TIS11B RBD fused to the N-terminus of SUMO10-SIM5) fusion protein, two overlapping fragments were PCR amplified. Fragment 1, mCherry, was PCR amplified from the mCherry-TIS11B construct with primers TIS-SUMO-SIM 1F and TIS-SUMO-SIM 1R. Fragment 2, the RBD (aa 114 to 181) of TIS11B, was PCR from the mCherry-TIS11B construct with primers TIS-SUMO-SIM 2F and TIS-SUMO-SIM 2R. A ligation PCR was performed to generate mCherry-TIS11B RBD with primers SUMO-SIM 1F and TIS-SUMO-SIM 2R. mCherry-TIS11B RBD was cloned into the SUMO10-SIM5 construct with AgeI and BsrGI restriction sites.

For the FUS-TIS (FUS-IDR fused to the N-terminus of TIS11B RBD) fusion protein, two overlapping fragments were PCR amplified. Fragment 1: the IDR (aa 1 to 214) of FUS, was PCR amplified from the pcDNA-puro-eGFP-FUS-3′UTR vector with primers FUS-TIS 1F and FUS-TIS 1R. Fragment 2: the RBD (aa 114 to 181) of TIS11B was PCR amplified with primers FUS-TIS 2F and FUS-TIS 2R. A final ligation PCR was performed to ligate two PCR fragments to the full-length FUS-TIS chimera with primers FUS-TIS 1F and FUS-TIS 2R. The full-length FUS-TIS was cloned into pcDNA3.1-puro-mGFP vector with BsrGI and EcoRI restriction sites.

### Transfections

Lipofectamine 2000 (Invitrogen) was used for all transfections.

### RNA oligonucleotide pulldown

To examine the RNA-binding activity of TIS11B WT and TIS11B RBD mutant, RNA oligonucleotide pulldown was performed as described previously (*9*). A 3′-biotinylated RNA oligonucleotide of the *TNFα* ARE-1 (AU-rich element) was purchased from Dharmacon. mCherry-tagged constructs were transfected into HeLa cells with or without 3′-biotinylated RNA oligonucleotides. Twenty-four hours after transfection, HeLa cells were lysed with 200 μl ice-cold NP-40 lysis buffer (25 mM Tris-HCl pH 7.5, 150 mM NaCl, 1% NP-40, 1 mM EDTA) for 30 min. Then, cell lysates were spun down at 20,000 g for 10 min at 4°C. The supernatant was transferred to a pre-cooled tube and diluted with 300 μl ice-cold dilution buffer (10 mM Tris-HCl pH 7.5, 150 mM NaCl, 0.5 mM EDTA). Streptavidin C1 beads (Invitrogen) were added to each tube and rotated for 1 hour at 4°C. Beads were washed three times with wash buffer (10 mM Tris-HCl pH 7.5, 150 mM NaCl, 0.5 mM EDTA). Lastly, 2x Laemmli sample buffer was added to the beads, boiled at 95°C for 10 min and cooled on ice before loading on SDS page gels. This was followed by Western blotting.

### Western blot

Western blots were performed as described previously (*9*). Imaging was captured on the Odyssey CLx imaging system (Li-Cor). The antibodies used are mouse anti-α-TUBULIN (Sigma-Aldrich, T9026), Mouse anti-mCherry (Abcam, ab125096), and Rabbit anti-HuR (Millipore, 07-1735).

### Recombinant protein purification

mGFP-FUS-TIS was cloned into the bacterial expression vector pET28a, which was a gift from the lab of Dirk Remus (MSKCC). At the N-terminus of mGFP, we added a 6xHis-MBP tag, followed by a Tev protease cleavage site. At the C-terminus of TIS11B, we added a Strep-Tag II (SAWSHPQFEK). The 6xHis-MBP tag was PCR amplified from pDZ2087 construct (Addgene, #92414) with primers MBP F and MBP R. Full-length 6xHis-MBP was cloned into pET28a backbone with XbaI and EcoRI restriction sites. mGFP-FUS-TIS-Strep-tag II was PCR amplified from pcDNA-mGFP-FUS-TIS construct with primers mGFP F and TIS RBD-Strep-tag R. The Strep-tag II sequence was incorporated into primer TIS RBD-Strep-tag R. Full-length mGFP-FUS-TIS-Strep-tag II was cloned into pET28a-6xHis-MBP backbone with NheI and EcoRI restriction sites.

To purify high-quality FUS-TIS protein, we used three steps of purification. Step 1: His-Ni purification; Step 2: Strep-Tag II purification; Step 3: size exclusion chromatography. pET28a-6xHis-MBP-Tev cleavage site-mGFP-FUS-TIS-Strep-tag II was transformed into BL21 E.coli (New England Biolabs). Two fresh colonies were cultivated overnight in 2 × 50 ml SOB medium at 37 °C, and then 4 × 25 ml bacteria were transferred to 4 × 1 L SOB medium to grow at 37 °C until OD600 reached 0.6. Bacteria were then kept at a 4 °C cold room until 18:00. Protein expression was induced by adding of 1 mM IPTG, and bacteria were incubated at 16 °C overnight.

Bacteria were centrifuged at 6,000 g for 10 min, and the pellet was resuspended in 100 ml cold lysis buffer. High salt lysis buffer (1 M NaCl, 25 mM Tris-Cl, pH 8.0, 20 mM imidazole, 1 mM DTT, 1x PMSF) was used to remove nucleic acid contamination. Bacteria were sonicated on ice for 60 min with on/off interval of 1 and 2 seconds. The lysate was centrifuged at 13,000 g for 30 min.

6 ml Ni-NTA (QIAGEN) was washed with 5 column volumes of wash buffer 1 (150 mM NaCl, 25 mM Tris-Cl, pH 8.0, and 1 mM DTT). After centrifugation, the supernatant of the bacteria lysate was transferred into new 50 ml Falcon tubes and incubated with Ni-NTA (Qiagen) at 4 °C for 30 min. Then the sample was transferred into three gravity columns and washed respectively with 40 ml wash buffer 2 (1 M NaCl, 25 mM Tris-Cl, pH 8.0, 20 mM imidazole, and 1 mM DTT), followed with 10 ml wash buffer 3 (600 mM NaCl, 25 mM Tris-Cl, pH 8.0, 20 mM imidazole, and 1 mM DTT). Then, the sample was eluted with 30 ml elution buffer 1 (600 mM NaCl, 25 mM Tris-Cl, pH 8.0, 200 mM imidazole, and 1 mM DTT).

After Ni-NTA purification, the eluted sample was transferred to a 5 ml StrepTrap column (GE Healthcare, Cat. No. 28907547), which was pre-equilibrated with elution buffer 2 (600 mM NaCl, 20 mM Tris-HCl, pH 7.4, and 1 mM DTT) using the AKTA Purifier system (GE Healthcare). The target protein was eluted with 20 ml elution buffer 3 (600 mM NaCl, 20 mM Tris-HCl, pH 7.4, 2.5 mM desthiobiotin (Sigma-Aldrich), and 1 mM DTT).

The eluted protein was concentrated using Amicon Ultra-centrifugal filters-50K (Millipore). The concentrated sample was further purified by gel filtration on HiLoad 16/600 Superdex200 column (GE Healthcare) in elution buffer 2 (600 mM NaCl, 20 mM Tris-HCl, pH 7.4, and 1 mM DTT) using the AKTA Purifier system (GE Healthcare).

The fractions representing the monomeric protein were collected and concentrated with Amicon Ultra-centrifugal filters-50K (Millipore). The quality of the final protein product was examined by SDS PAGE. The protein concentration was measured by Bradford assay (Biorad). The protein was aliquoted into PCR tubes and flash-frozen in liquid nitrogen and then stored at -80 °C.

### *In vitro* transcription of RNA

All RNAs were *in vitro* transcribed using the T7 MEGAscript kit (Ambion by life technologies). All DNA templates used for in vitro transcription were PCR amplified and purified with a gel extraction kit (QIAGEN). The T7 promoter (TAATACGACTCACTATAGGG) was incorporated into the forward primers used to amplify the DNA templates.

The DNA sequences from the full-length 3′UTRs of *CD47, ELAVL1, CD274*, and *FUS* were PCR amplified from pcDNA-eGFP-CD47-3′UTR, eGFP-ELAVL1-3′UTR, eGFP-CD274-3′UTR, and eGFP-FUS-3′UTR constructs. The DNA sequences of the 3′UTRs *of CD44, VSIG10, IL10, TNFSF11, GPR39, TLR8, GPR34, TNFAIP6, MYC, PLA2G4A, HEATR5B, PPP1R3F, DRD1, FAM72B, MCOLN2, TSPAN13, LHFPL6, FAM174A, VPS29, ADPGK, ASPN, CASP8, CLCA2, EOMES, ESCO1, GLYATL3, HNRNPH3, HOGA1, LPAR4, LRBA, LYPLAL1, ODF2, PRKDC, RHOA, SHQ1, SLC39A6, SLC5A9, SMIM3, SNTN, SOSTDC1, STBD1, TP53TG3*, and *TTC17* were PCR amplified from HeLa genomic DNA.

To generate the *TNFSF11* 3′UTR mutant carrying two 15-nt oligo insertions, two overlapping fragments were PCR amplified. Fragment 1: *TNFSF11* 3′UTR with oligo 1 using primers *TNFSF11 3′UTR T7 F* and *TNFSF11* mutant R1. Fragment 2: *TNFSF11* 3′UTR with oligo 1 and oligo 2 using primers *TNFSF11* mutant F2 and *TNFSF11* mutant R2. The inserted 15-nt oligo 2 sequence was incorporated into the primer *TNFSF11* mutant R2. A final ligation PCR was performed to ligate two PCR fragments to generate the full-length *TNFSF11* 3′UTR mutant with primers *TNFSF11 3′UTR T7 F* and *TNFSF11* mutant R2.

RNAs with exogenous dimerization elements (D1a, crRNA, D1b, tracrRNA, D2, HIV dimerization motif) were generated as follows. For D1a-*TLR8*-D2, two rounds of PCR were performed. Round 1: primers D1a-TLR8-D2 1F and D1a-TLR8-D2 1R; Round 2: primers D1a-TLR8-D2 2F and D2 R. For D1b-*MYC*-D2, three rounds of PCR were performed. Round 1: primers D1b-MYC-D2 1F and MYC 3′UTR R; Round 2: primers D1b-MYC-D2 2F and D1b-MYC-D2 2R; Round 3: primers D1b-MYC-D2 3F and D2 R.

*In vitro* transcription was performed in a 20 μl volume according to the manufacturer’s guidelines. To generate Cy3 or Cy5 labeled RNA, 0.2 μl of 2.5 mM Cy3-UTP or Cy5-UTP (Enzo Life Sciences) was added into the *in vitro* transcription reaction.

The transcription reaction was incubated 3 hours at 37 °C in a PCR machine. All transcribed RNAs were digested with DNase for 30 min at 37°C, then precipitated with LiCl for 4 hours to overnight at - 20 °C. RNAs were centrifuged at 13,000 for 15 min, and the RNA pellets were washed with 70% ethanol three times. RNAs were dissolved in Nuclease-free water and stored at - 20 °C. The concentration of RNAs was measured by Nanodrop one.

### *In vitro* phase separation assay

To allow phase separation, purified 6xHis-MBP-mGFP-FUS-TIS-Strep tag II protein stock was incubated with Tev protease for 1 hour at room temperature to cleave off the 6xHis-MBP tag. mGFP and Strep tag II were not cleaved off. All phase separation assays were performed in 20 μl phase separation buffer (150 mM NaCl, 200 μM ZnCl2, 25 mM Tris-Cl, pH 7.4, 1 mM DTT, 2.5% glycerol, 5% Dextran T500 (Pharmacosmos)). ZnCl2 was added as the RNA-binding domain of TIS11B has two zinc finger motifs. Only in the phase separation assay shown in Figure S2 ZnCl2 was omitted.

After 1 hour of Tev protease digestion, the FUS-TIS protein stock was diluted into the desired concentrations with protein stock buffer (600 mM NaCl, 25 mM Tris-Cl, pH 7.4, 1 mM DTT) and centrifuged at 13,000 g for 2 min to remove small protein aggregates. The supernatant was transferred into a new Eppendorf tube. The phase separation assay was mixed in PCR tubes. Dextran buffer and RNAs with desired concentrations were first mixed in PCR tubes, then FUS-TIS protein was added into the PCR tube and immediately mixed thoroughly. The final concentration of FUS-TIS and RNAs are indicated in the figures. The mixture (20 μl) was then transferred into a 384-well glass-bottom microplate (Greiner bio-one). The chambers of the microplate were pre-treated with 1 mg/ml BSA (NEB) for 30 min before aspirating the BSA. The microplate was kept in the dark at room temperature for two or 16 hours, followed by imaging of the condensates using confocal microscopy.

### Confocal microscopy

Confocal imaging was performed using ZEISS LSM 880 with Airyscan super-resolution mode. A Plan-Apochromat 63x/1.4 Oil objective (Zeiss) was used. For live cell imaging, HeLa cells were plated on 3.5 cm glass-bottom dishes (Cellvis) and transfected with the indicated constructs. Fourteen hours after transfection, cells were imaged in cell culture medium while incubating in a LiveCell imaging chamber (Zeiss) at 37°C and 5% CO2. Images were prepared with the commercial ZEN software black edition (Zeiss).

### Fluorescence recovery after photobleaching (FRAP)

FRAP experiments were performed with ZEISS LSM 880 confocal microscopy. A Plan-Apochromat 63x/1.4 Oil objective (Zeiss) was used. 10 μM mGFP-FUS-TIS was mixed with 50 nM *CD47* 3′UTR to induce granule network formation. Two hours after mix, an area of diameter = 1 μm was bleached with a 405 nm and 633 nm laser. GFP fluorescence signal was collected over time. The fluorescence intensity of the bleached area was obtained by ZEN software black edition (ZEISS). The prebleached fluorescence intensity was normalized to 1, and the signal after bleach was normalized to the prebleach level.

### RNA native agarose gel electrophoresis

RNA native agarose gel electrophoresis was performed as described previously with a few modifications (*19*). For sphere-forming and network-forming RNAs, RNAs were diluted into 5 μl buffer A (150 mM NaCl, 25 mM Tris-Cl, pH 7.4) to a final concentration of 5 μM. RNAs were incubated at 95 °C for 2 min in a PCR machine and then incubated on ice for 2 min. RNAs were kept at 37 °C for two hours. 1 μl native agarose gel loading buffer (6X stock: 60% glycerol, 10 mM Tris-Cl, pH 7.4, 0.03% bromophenol blue, and 0.03% xylene cyanol FF) was added into the RNA. A total of 1 μg RNA was loaded into the 1% agarose gel made with the TAE (Tris-acetate-EDTA) buffer for electrophoresis with TAE buffer.

For RNAs containing dimerization elements (*TLR8* 3′UTR, *MYC* 3′UTR, D1a-*TLR8*-D2, D1b-*MYC*-D2), each RNA was diluted into 5 μl buffer A (150 mM NaCl, 25 mM Tris-Cl, pH 7.4) to a final concentration of 2 μM. RNAs were incubated at 95 °C for 2 min in a PCR machine and then incubated on ice for 2 min. 2.5 μl *TLR8* 3′UTR or 2.5 μl *MYC* 3′UTR were each diluted with 2.5 μl buffer A. 2.5 μl *TLR8* 3′UTR and 2.5 μl *MYC* 3′UTR were mixed together. 2.5 μl D1a-TLR8-D2 or 2.5 μl D1b-MYC-D2 were each diluted with 2.5 μl buffer A. 2.5 μl D1a-TLR8-D2 and 2.5 μl D1b-MYC-D2 were mixed together. The final concentration of each RNA was 1 μM in 5 μl buffer A. RNAs were kept at 37 °C for two hours. 1 μl native agarose gel loading buffer was added to the RNA. A total of 1 μg RNA was loaded onto the 2% agarose gel made with the TBE (Tris-borate-EDTA) buffer for electrophoresis with TBE buffer.

### Calculation of mRNA concentration in HeLa cells

We estimated the concentration of a specific mRNA in mammalian cells is between 8 pM – 8 nM (9.5 pg/μl - 9.5 ng/μl) based on the following assumptions: (1) The volume of a HeLa cell is 2000 μm^3^, (2) the average length of an mRNA is 3500 nt which corresponds to an average molecular weight of 1155 kDa, and (3) there are between 10 – 10,000 copies of mRNA per cell (*14, 15*).

### Calculation of NED values

The ensemble diversity of 3′UTR sequences was calculated using the RNAfold software (version: 2.4.14; command line: RNAfold --MEA -d2 -p --infile=<RNA_sequences.fasta> -- outfile=<RNA_sequences.RNAfold.summary>) (*17, 30*). Only 3′UTRs with a length < 7500 nucleotides can be analyzed by RNAfold. As the values for ensemble diversity depend on the sequence length, we calculated the ‘normalized ensemble diversity’ (NED) by dividing the value of ensemble diversity by the length of the 3′UTR in nucleotides. All values are listed in table S2. The cut-offs for high and low NED values were determined empirically. Using our experimental dataset of NED values with correctly predicted ability for sphere- or network formation (*N* = 36), only 1/19 3′UTRs that induced network formation had a NED value < 0.29. Conversely, only 2/17 3′UTRs that induced sphere formation had NED values > 0.27. Therefore, high NED values are considered as ≥ 0.29 and correlate with the presence of large IDRs in 3′UTRs and with formation of mesh-like condensates, whereas low NED values are considered ≤ 0.27 and correlate with the lack of large IDRs and with formation of sphere-like condensates.

### AU-rich elements and nucleotide content

For counting of AU-rich elements, we only considered the canonical sequence AUUUA. We counted the number of AU-rich elements in annotated 3′UTRs of mRNAs expressed in HeLa cells (*31*). The 3′UTR length is the full-length 3′UTR length obtained from Refseq. The nucleotide content of the 3′UTRs was calculated from the same dataset. All values are listed in table S2.

### Protein size and protein IDRs

Protein sequences and protein sizes were obtained from uniprot. The IUPRED2A software (https://iupred2a.elte.hu/) (*32, 33*) was used to identify disordered protein regions (command line: python iupred2a.py <protein.fasta> long > <iupred2.output>). We used our own script to extract disordered regions. A region was called disordered if 30 consecutive amino acids reached an iupred2 score > 0.5. All amino acids within these regions were considered as disordered. All values are listed in table S2.

### Statistical methods

For all pair-wise comparisons a two-sided Mann-Whitney test was performed. For comparisons containing more than two groups, a Kruskal-Wallis test was performed. The Pearson’s correlation coefficient is reported.

