## Supplementary figures for "Unstructured mRNAs form multivalent RNA-RNA interactions to generate TIS granule networks"

Ma et al. Figure S1

**A**

```
                        RBD WT     RYKTELCRPFEENGACKYGDKCQFAHGIHELRSLTRHPKYKTELCRTFHTIGFCPYGPRCHFIHNAEE
RBD mutant CC  (C135H/C173H)      RYKTELCRPFEENGACKYGDKHQFAHGIHELRSLTRHPKYKTELCRTFHTIGFCPYGPRHHFIHNAEE
RBD mutant FF  (F137N/F175N)      RYKTELCRPFEENGACKYGDKCQNAHGIHELRSLTRHPKYKTELCRTFHTIGFCPYGPRCHNIHNAEE
RBD mutant KK  (K116L/K154L)      RYLTELCRPFEENGACKYGDKCQFAHGIHELRSLTRHPKYLTELCRTFHTIGFCPYGPRCHFIHNAEE
RBD mutant RK  (R114L/K152L)      LYKTELCRPFEENGACKYGDKCQFAHGIHELRSLTRHPLYKTELCRTFHTIGFCPYGPRCHFIHNAEE
```

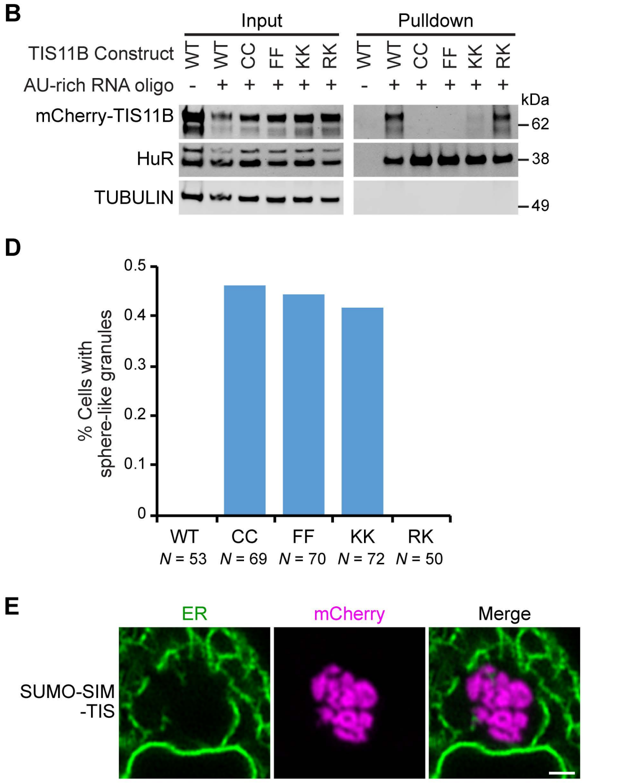


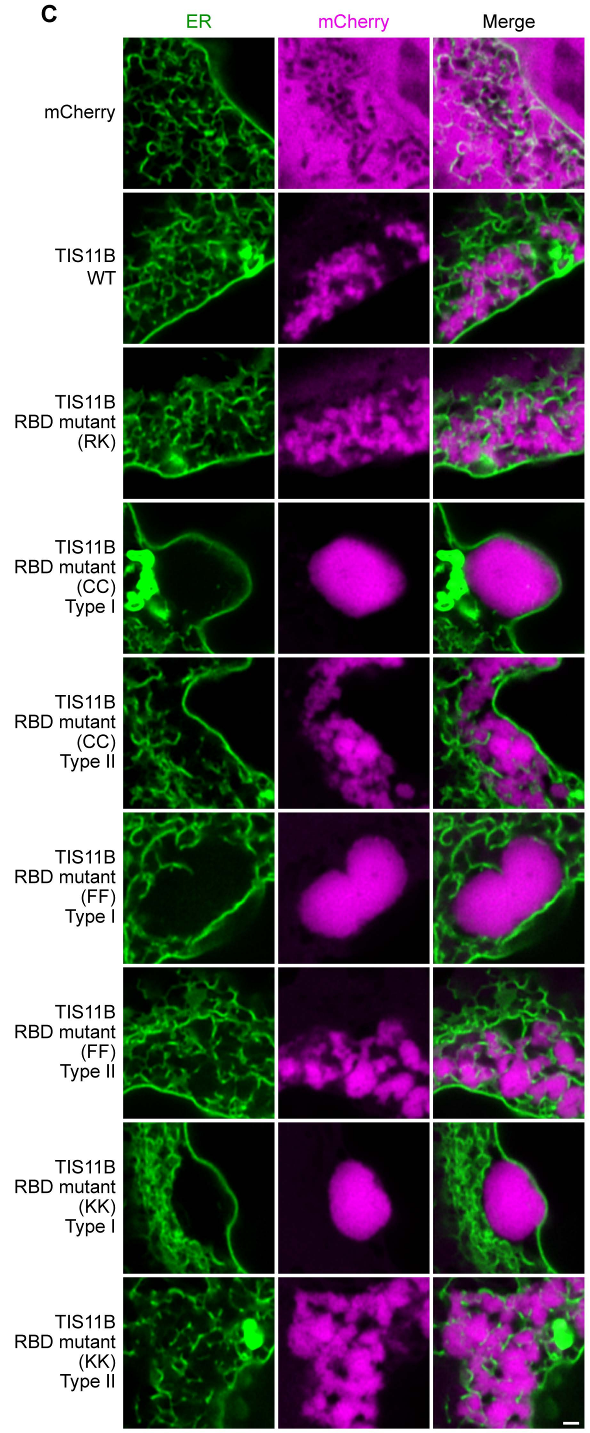


**Figure S1. Mutation of the TIS11B RNA-binding domain generates sphere-like granules in cells.**

(**A**) Shown is the amino acid sequence of the wild-type (WT) TIS11B RNA-binding domain (RBD) together with the introduced point mutations (red).

(**B**) Western blot showing RNA oligonucleotide pulldown after co-transfection of the indicated mCherry-tagged TIS11B constructs and an AU-rich RNA oligo (*TNFα* AU-rich element) into HeLa cells. RNA-binding is disrupted in CC, FF, and KK mutants. HuR also binds to the AU-rich element. Tubulin was used as loading control. 2.5% of the input was loaded.

(**C**) Confocal live cell imaging of HeLa cells after transfection of the indicated constructs described in (A). GFP-SEC61B was co-transfected to visualize the ER. TIS11B containing a mutated RBD (CC, FF, or KK) form two types of granules in cells. Type I are sphere-like granules, whereas type II form an intermediate between sphere-like and network-like granules.

(**D**) Quantification of the data shown in (C). Shown is the fraction of cells with sphere-like granules formed by the indicated TIS11B proteins.

(**E**) Same as (C), but after transfection of SUMO-SIM-TIS chimera.

Ma et al. Figure S2

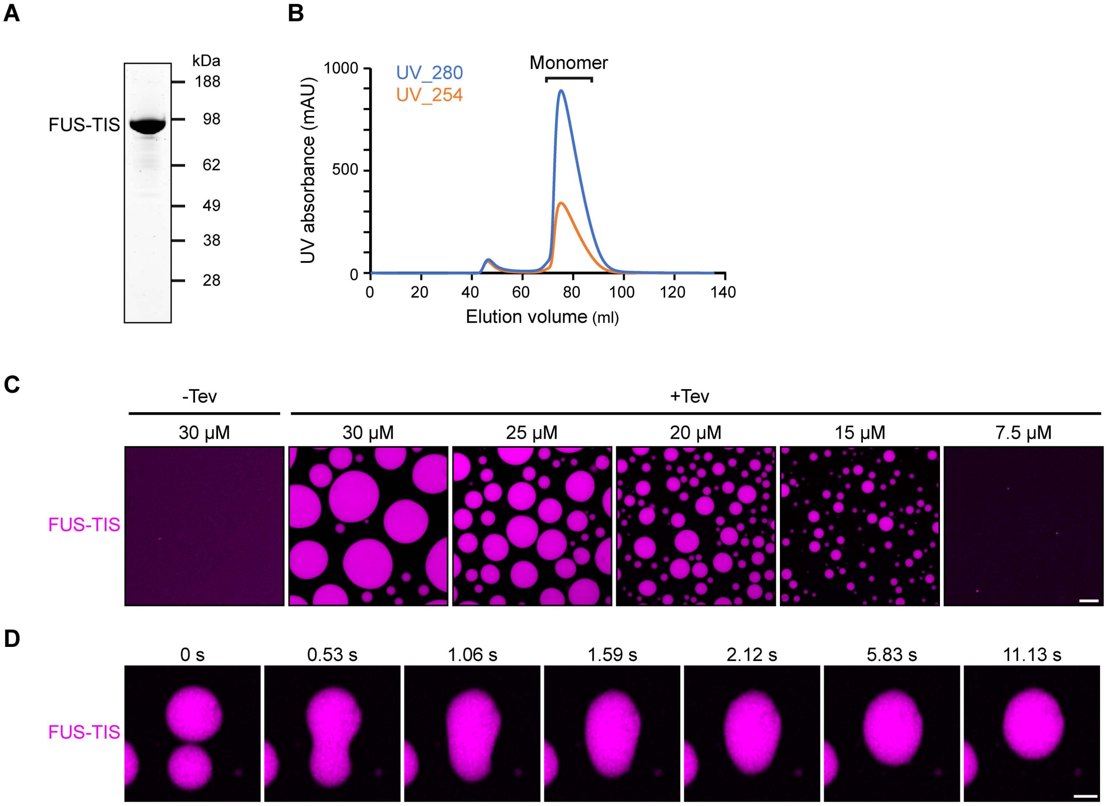


**Figure S2. Purified FUS-TIS protein phase separates *in vitro*.**

(**A**) SDS-PAGE of the purified FUS-TIS protein. FUS-TIS was tagged with His-MBP-mGFP at the N-terminus and with Strep-tag II at the C-terminus and purified from E. coli. mGFP, monomeric GFP.

(**B**) Size exclusion chromatography of the FUS-TIS protein.

(**C**) Phase separation of purified FUS-TIS protein at the indicated concentrations was induced by reducing the salt concentration from 600 mM to 150 mM through dilution; shown is a 2 hour time point. The His-MBP tag was cleaved off before the phase separation experiment using TEV protease. Scale bar, 10 μm.

(**D**) Snapshots show a fusion event of FUS-TIS condensates at 2 hours.

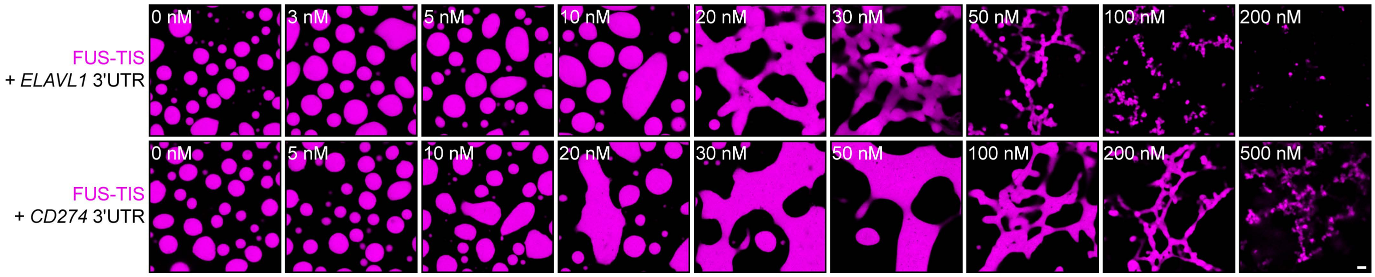


**Figure S3. Specific RNAs induce granule network formation *in vitro*.**

Representative confocal images of phase separation experiments using purified mGFP-FUS-TIS (10 µM) in the absence or presence of the indicated *in vitro* transcribed RNAs after 16 hours of incubation. Scale bar, 2 µm.

Ma et al. Figure S4

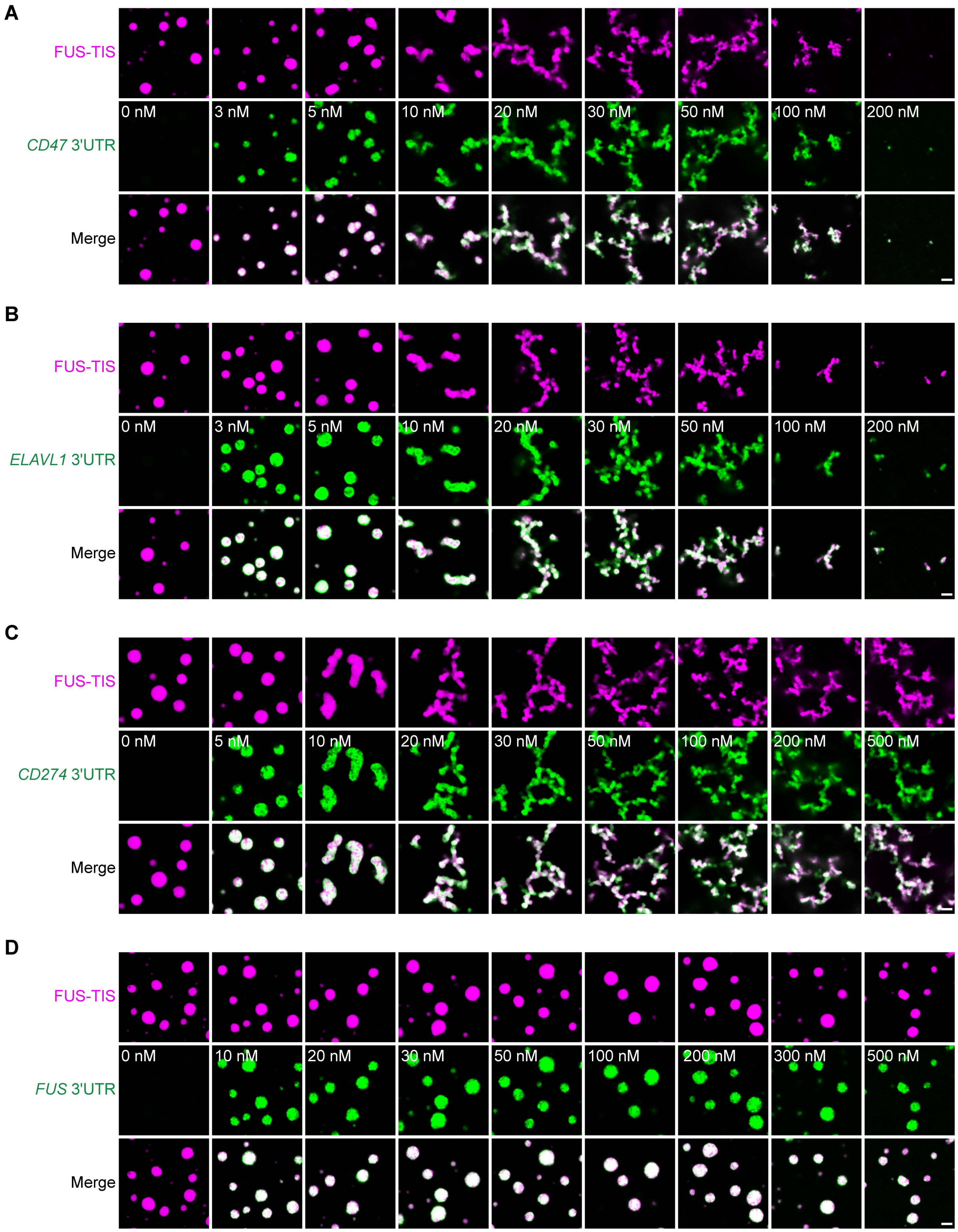


**Figure S4. Specific RNAs induce granule network formation *in vitro*.**

(**A** to **D**) Representative confocal images of phase separation experiments using purified mGFP-FUS-TIS (10 μM) in the absence or presence of the indicated *in vitro* transcribed RNAs after 2 hours of incubation. Scale bar, 2 μm.

Ma et al. Figure S5

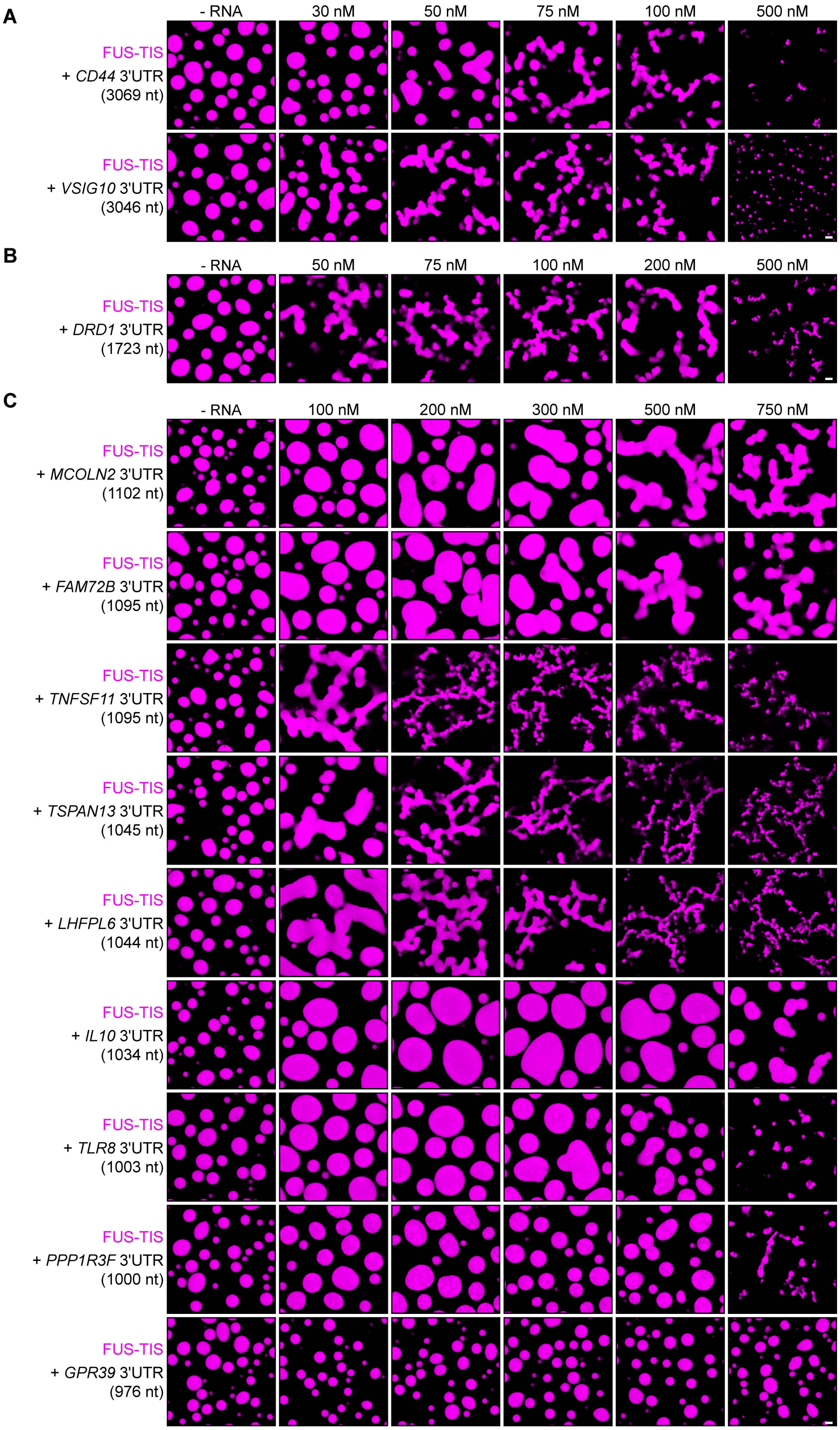


**Figure S5. Specific RNAs induce granule network formation *in vitro*.**

Representative confocal images of phase separation experiments using purified mGFP-FUS-TIS (10 μM) in the absence or presence of the indicated *in vitro* transcribed RNAs after 16 hours of incubation. Scale bar, 2 μm.

(**A**) RNAs with a length of approximately 3000 nt are shown.

(**B**) RNAs with a length between 1500 - 2000 nt are shown.

(**C**) RNAs with a length of approximately 1000 nt are shown.

Ma et al. Figure S6

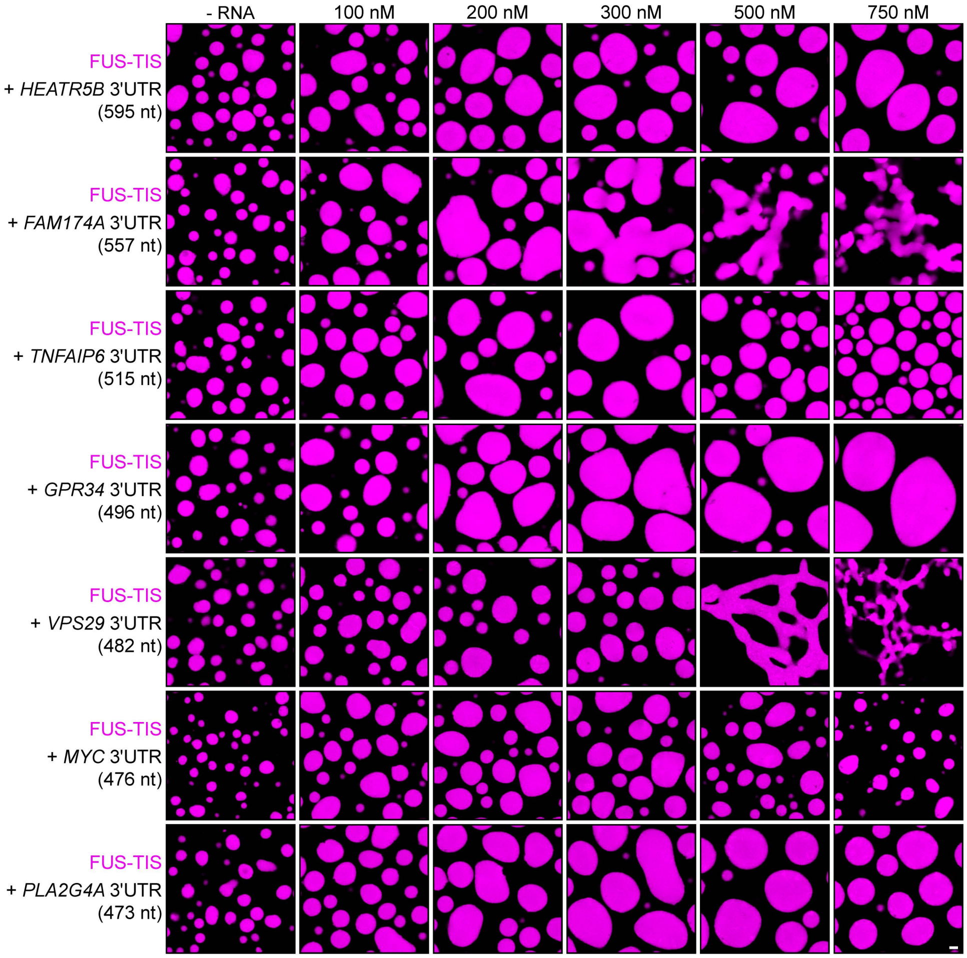


**Figure S6. Specific RNAs induce granule network formation *in vitro*.**

Representative confocal images of phase separation experiments using purified mGFP-FUS-TIS (10 μM) in the absence or presence of the indicated *in vitro* transcribed RNAs after 16 hours of incubation. Scale bar, 2 μm. RNAs with a length of approximately 500 nt are shown.

Ma et al. Figure S7

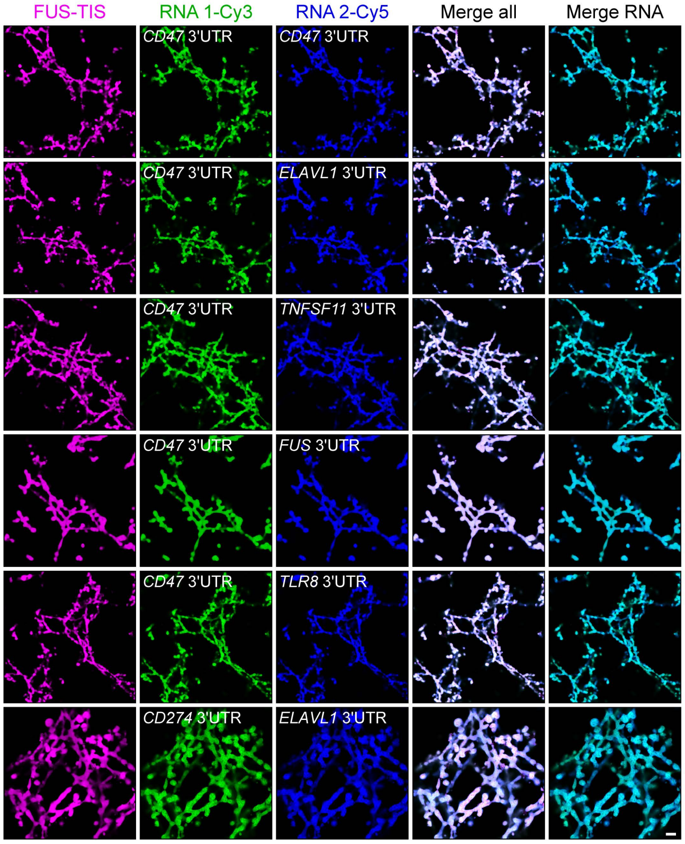


**Figure S7. RNAs co-localize in mesh-like condensates.** Representative confocal images of phase separation experiments using purified mGFP-FUS-TIS (10 μM) in the presence of two different *in vitro* transcribed RNAs that were labeled with Cy3 or Cy5 fluorescent dye, respectively. Images were taken after 16 hours of incubation. Scale bar, 5 μm.

Ma et al. Figure S8

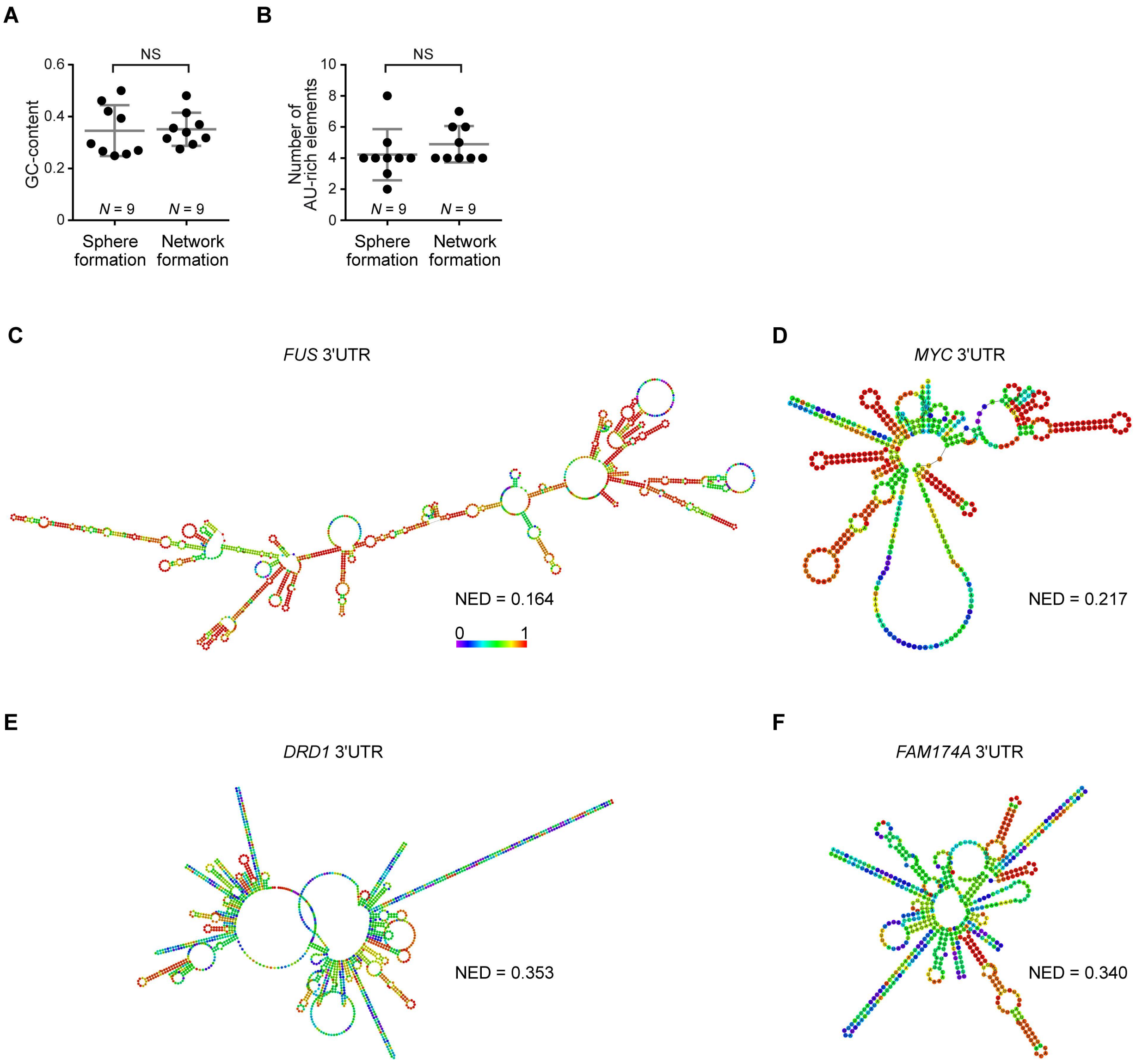


**Figure S8. Network-forming RNAs have weak local secondary structures.**

(**A**) Distribution of GC-content of sphere- and network-forming RNAs. See also table S1. Mann-Whitney test, Z=0.566, *P* = 0.605, NS, not significant.

(**B**) Number of AU-rich elements in sphere- and network-forming RNAs. See also table S1. Mann-Whitney test, Z=0.190, *P* =0.258, NS.

(**C** and **D**) Centroid RNA secondary structure of two sphere-forming 3′UTR predicted by RNAfold. The color code represents base-pairing probability. See also the URLs in table S1.

(**E** and **F**) As in (C), but structure is shown for two network-forming 3′UTRs.

Ma et al. Figure S9

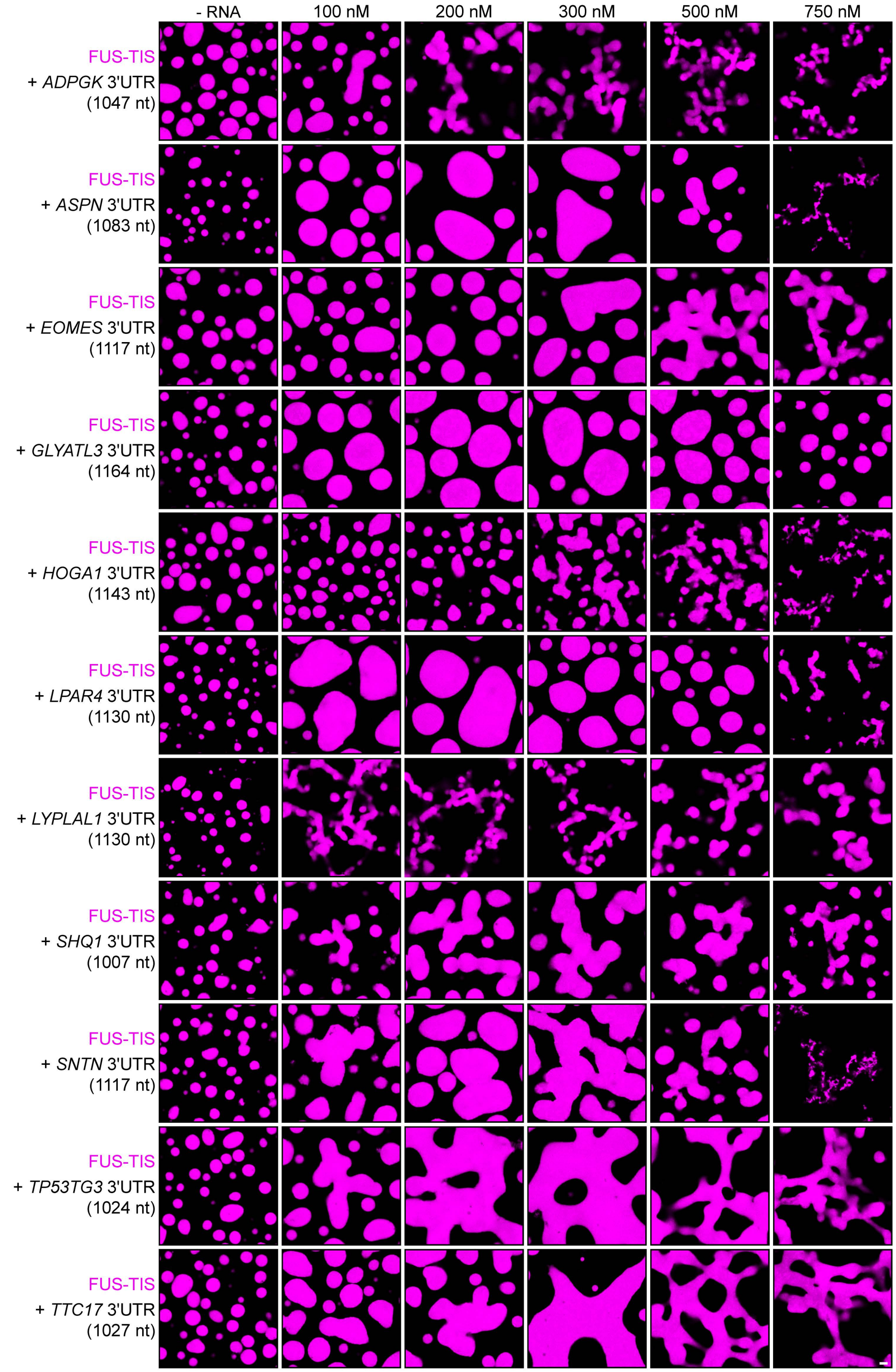


**Figure S9. Ten out of eleven RNAs with a high NED value were predicted correctly to induce network formation.** Representative confocal images of phase separation experiments using purified mGFP-FUS-TIS (10 μM) in the absence or presence of the indicated *in vitro* transcribed RNAs after 16 hours of incubation. Scale bar, 2 μm. The length of all RNAs is approximately 1000 nt. The minimum concentration for induction of network formation varies. The *GLYATL3* 3′UTR is unable to induce network formation even at high concentration. See also table S1.

Ma et al. Figure S10

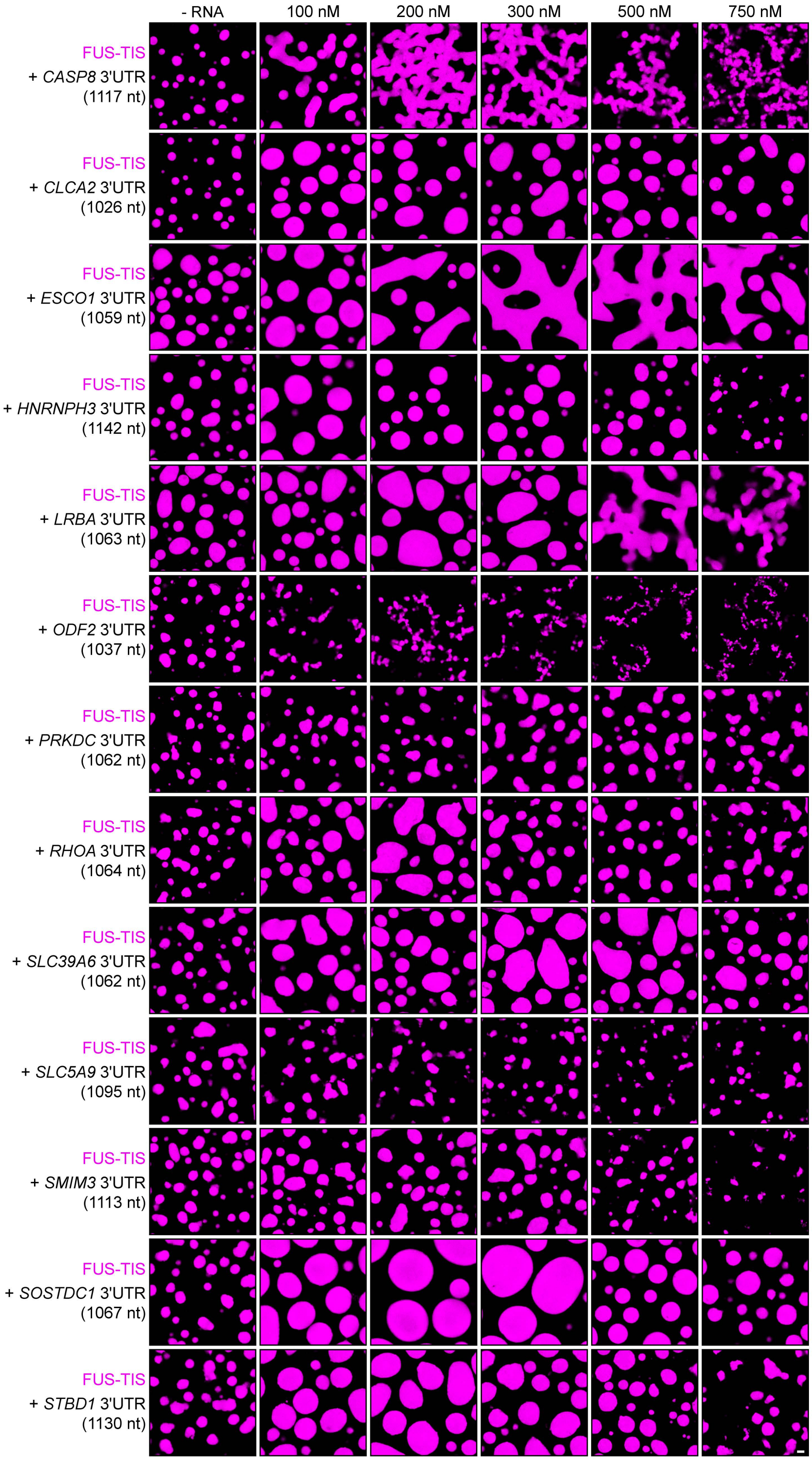


**Figure S10. Nine out of thirteen RNAs with a low NED value were predicted correctly to induce sphere-like condensate formation.**
Representative confocal images of phase separation experiments using purified mGFP-FUS-TIS (10 µM) in the absence or presence of the indicated *in vitro* transcribed RNAs after 16 hours of incubation. Scale bar, 2 µm. The length of all RNAs is approximately 1000 nt. See also table S1.

Ma et al. Figure S11

*TNFSF11* 3'UTR - mutant

GCCCCAGUUUUUGGAGUGUUAUGUAUUUCCUGGAUGUUUGGAAACAUUUUUUAAAACAAGCCAAGAAAGAUGUAUAUAGGUGUGUGAGACUACUAAGAGGCAUGGCCCCAA
CGGUACACGACUCAGUAUCCAUGCUCUUGACCUUGUAGAGAACACGCGUAUUUACAGCCAGUGGGAGAUGUUAGACUCAUGGUGUGUUACACAAUGGUUUUUAAAUUUUGU
AAUGAAUUCCUAGAAUUAAACCAGAUUGGAGCAAUUACGGGGUGACCUUAUGAGAAACUGCAUGUGGGCUAUGGGAGGGGUUGGUCCCUGGUCAUGUGCCCCUUCGCAGCU
GAAGUGGAGAGGGUGUCAUCUAGCGCAAUUGAAGGAUCAUCUGAAGGGGCAAAUUCUUUUGAAUUGUUACAUCAUGCUGGAACCUGCAAAAAAUACUUUUUCUAAUGAGGA
GAGAAAAUAUAUGUAUUUUUAUAUAAUAUCUAAAGUUAUAUUUCAGAUGUAAUGUUUUCUUUGCAAAGUAUUGUAAAUUAUAUUUGUGCUAUAGUAUUUGAUUCAAAAUAU
UUAAAAAUGUCUUGCUGUUGACAUAUUUAAUGUUUUAAAUGUACAGACAUAUUUAACUGGUGCACUUUGUCAGUUUCUCAUAAGGAAAUUCCCUGGGGAAAACUUGCAGCU
AAGGAGGGGAAAAAAAUGUUGUUUCCUAAUAUCAAAUGCAGUAUAUUUCUUCGUUCUUUUUAAGUUAAUAGAUUUUUUCAGACUUGUCAAGCCUGUGCAAAAAAAUUAAAA
UGGAUGCCUUGAAUAAUAAGCAGGAUGUUGGCCACCAGGUGCCUUUCAAAUUUAGAAACUAAUUGACUUUAGAAAGCUGACAUUGCCAAAAAGGAUACAUAAUGGGCCACU
GAAAUCUGUCAAGAGUAGUUAUAUAAUUGUUGAACAGGUGUUUUUCCACAAGUGCCGCAAAUUGUACCUUUUUUGUUUUUUCAAAAUAGAAAAGUUAUUAGUGGUUUAUCA
GCAAAAAAGUCCAAUUUUAAUUUAGUAAAUGUUAUCUUAUACUGUACAAUAAAAACAUUGCCUUUGAAUGUUAAUUUUUUGGUACAAAAAUAAAUUUAUAUGAAAACCUGC
GCAGGUUUUCAUAUA

**Figure S11. Nucleotide sequence of the *TNFSF11* 3′UTR mutant.**

Two 15-nt oligonucleotides (red and magenta bars) which are complementary to upstream sequences (red and magenta fonts) were added into the *TNFSF11* 3′UTR. This reduces the unstructured regions and increases the local secondary structure (Fig. 3E).

Ma et al. Figure S12

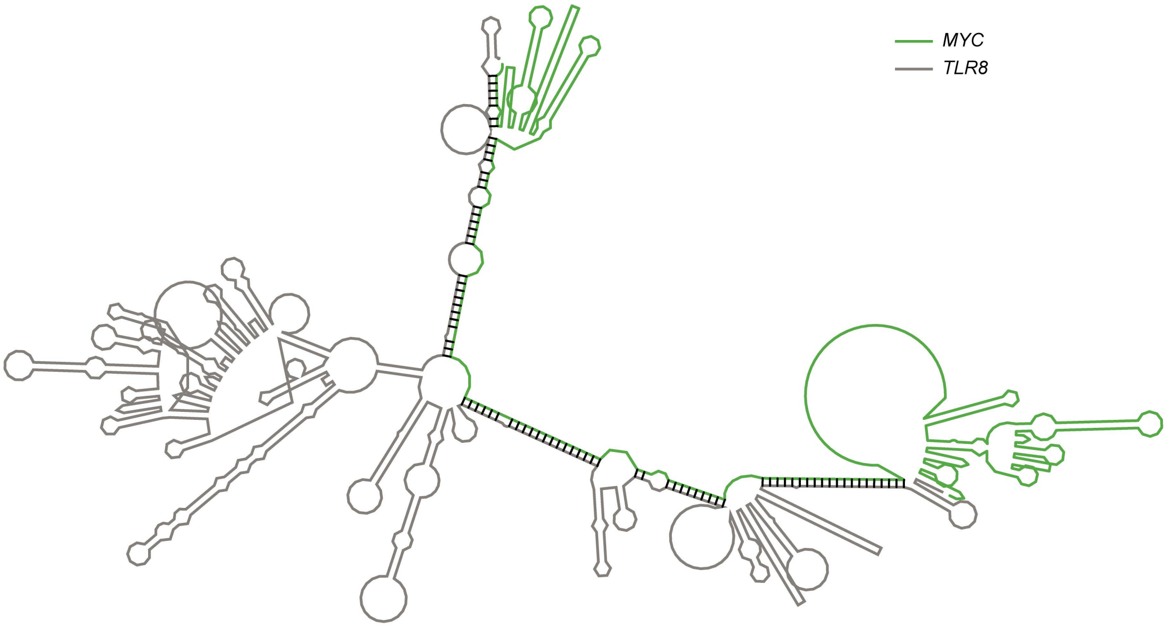


**Figure S12.** ***TLR8*** **3′UTR and** ***MYC*** **3′UTR are predicted to dimerize.**

The RNA secondary structure of the *TLR8* 3′UTR together with the *MYC* 3′UTR was predicted by RNAfold. *TLR8* 3′UTR is shown in grey, and *MYC* 3′UTR is shown in green. The secondary structure prediction shows extensive base pairing between the two RNAs, indicated by the black marks. The *MYC* 3′UTR was added directly downstream of the *TLR8* 3′UTR for structure prediction. Although both RNAs are able to base pair, they are unable to form complex RNA network.

Ma et al. Figure S13

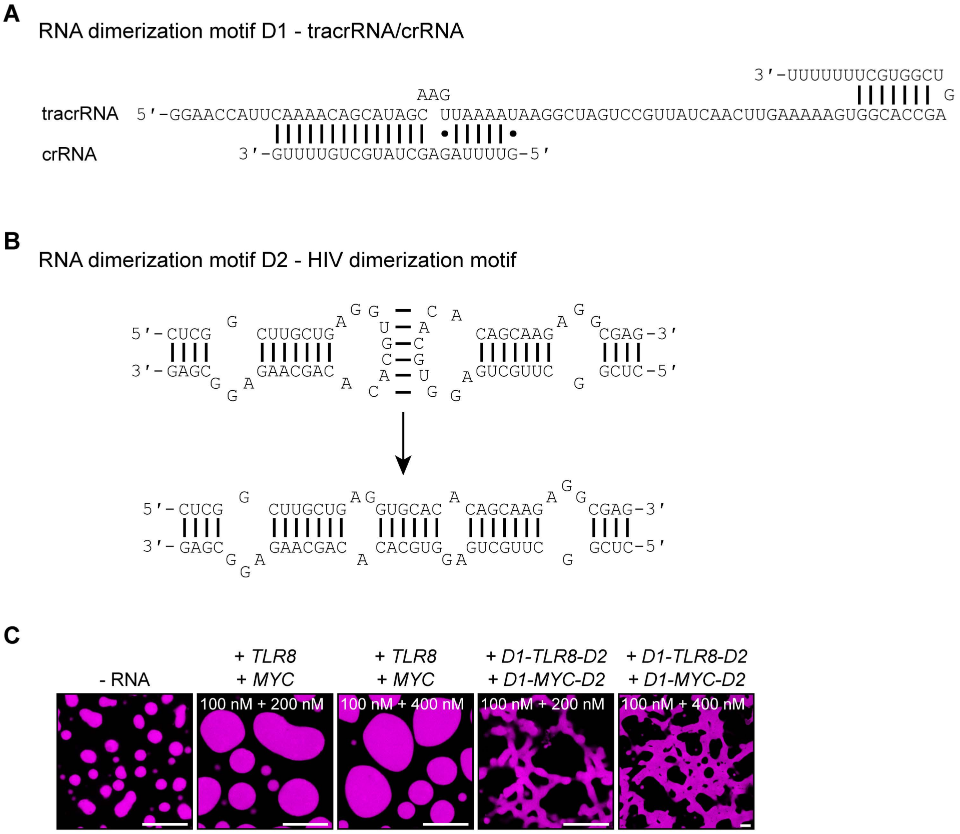


**Figure S13. Schematic of the RNA dimerization motifs.**

(**A**) Shown is the sequence and base-pairing of the directional RNA dimerization motif D1 which was obtained from the CRISPR/Cas9 and represents tracrRNA/crRNA. The two components form a heterologous RNA-RNA interaction.

(**B**) Shown is the sequence and base-pairing of the RNA dimerization motif D2 which is the HIV dimerization motif. It forms a homologous RNA-RNA interaction.

(**C**) Representative images of phase separation experiments using purified mGFP-FUS-TIS (10 μM) in the presence of the indicated *in vitro* transcribed RNAs after 16 hours of incubation. Scale bars, 10 μm.

Ma et al., Figure S14

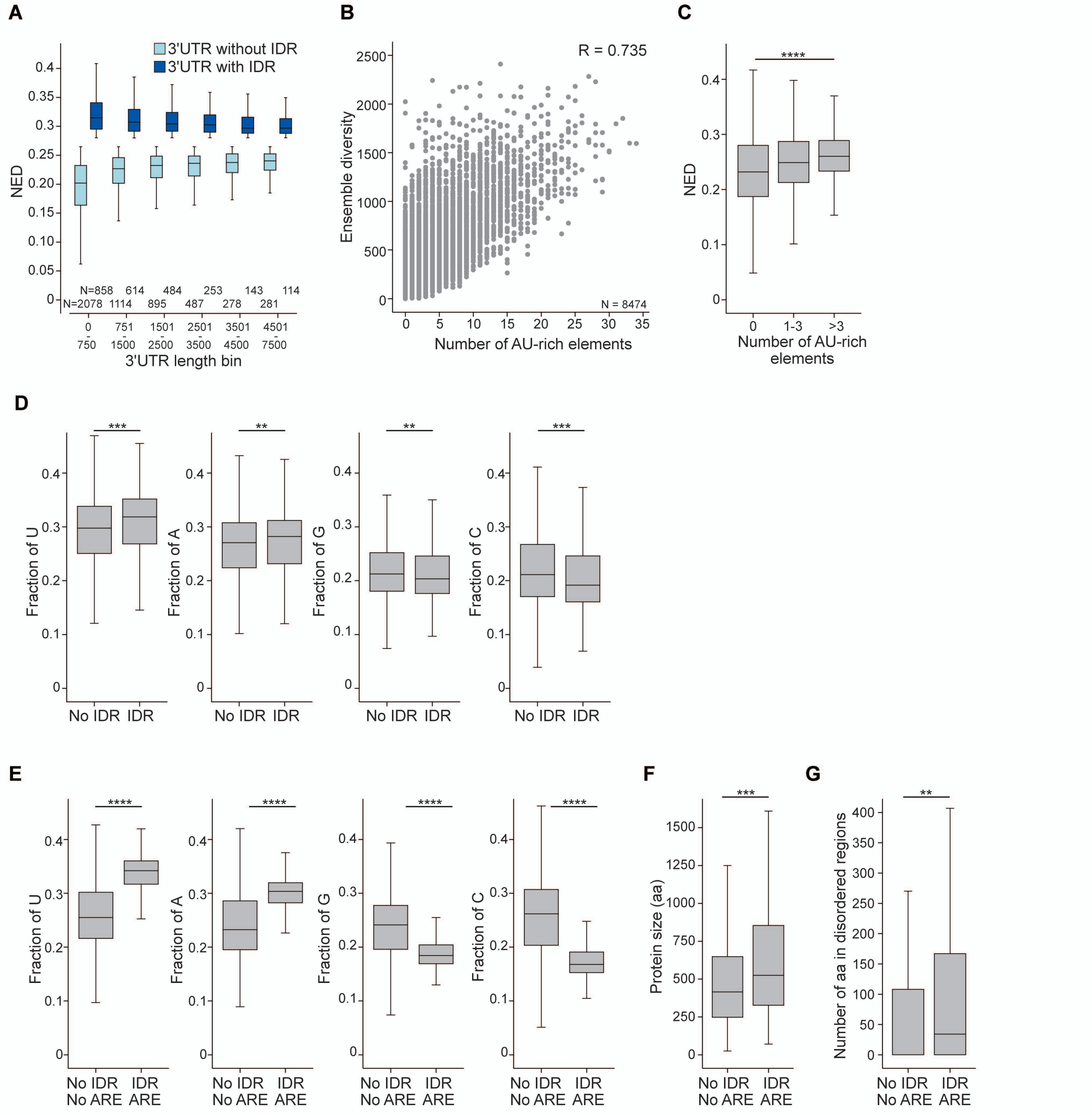


**Figure S14. mRNAs with IDRs are enriched in AU-rich elements.**

(**A**) The 3′UTRs of mRNAs expressed in HeLa cells were separated into 3′UTRs with high (NED ≥ 0.29) and low NED (NED ≤ 0.27) values. NED is a length-normalized ensemble diversity parameter. Shown is that NED values are largely independent of 3′UTR length after normalization. (**B**) Ensemble diversity correlates with the number of AU-rich elements in 3′UTRs. The Pearson's correlation coefficient is shown. *P* = 0. (**C**) The number of AU-rich elements correlates with NED. 3′UTRs without AU-rich elements, *N* = 2504, 3′UTRs with 1-3 AU-rich elements, *N* = 3081, 3′UTRs with more than three AU-rich elements, *N* = 2889 are shown. Kruskal-Wallis test: X2= 315, *P* = 4x10-69. (**D**) 3′UTRs with IDRs (N = 2466) have higher U-content compared with 3′UTRs without IDRs (*N* = 5113), Mann-Whitney test: Z=-11.2, *P* = 1x10-29. They also have a higher A-content (Z=-5.1, 1x10-7), but lower G-content (Z=-4.6, 1x10-6) and lower C-content (Z=-10.5, 1x10-26). (**E**) 3′UTRs with IDRs and more than three AU-rich elements (*N* = 939) have higher U-content compared with 3′UTRs without IDRs and without AU-rich elements (*N* = 1657), Mann-Whitney test: Z=-29.5, *P* = 1x10-191. They also have a higher A-content (Z=-13.3, 1x10-119), but lower G-content (Z=-22.1, 1x10-107) and lower C-content (Z=-27.8, 1x10-170). (**F**) As in (E), but shown is protein size in amino acids (aa). Mann Whitney test, Z=-28.9, 1x10-19. (**G**) As in (E), but shown is the number of aa in disordered regions. Only regions with at least 30 consecutive disordered aa were counted. Mann-Whitney test, Z=-4.9, 1x10-7.
